# Histone H3K27me3 demethylases regulate human Th17 cell development and effector functions by impacting on metabolism

**DOI:** 10.1101/820845

**Authors:** Adam P Cribbs, Stefan Terlecki-Zaniewicz, Martin Philpott, Jeroen Baardman, David Ahern, Morten Lindow, Susanna Obad, Henrik Oerum, Brante Sampey, Palwinder K Mander, Henry Penn, Paul Wordsworth, Paul Bowness, Rab K Prinjha, Menno de Winther, Marc Feldmann, Udo Oppermann

## Abstract

T helper (T_h_) cells are CD4+ effector T cells that play an instrumental role in immunity by shaping the inflammatory cytokine environment in a variety of physiological and pathological situations. Using a combined chemico-genetic approach we identify histone H3K27 demethylases KDM6A and KDM6B as central regulators of human Th subsets. The prototypic KDM6 inhibitor GSK-J4 increases genome-wide levels of the repressive H3K27me3 chromatin mark and leads to suppression of the key transcription factor RORγt during Th17 differentiation, whereas in mature Th17 cells an altered transcriptional program leads to a profound metabolic reprogramming with concomitant suppression of IL-17 cytokine levels and reduced proliferation. Single cell analysis reveals a specific shift from highly inflammatory cell subsets towards a resting state upon demethylase inhibition. The apparent root cause of the observed anti-inflammatory phenotype in stimulated Th17 cells is reduced expression of key metabolic transcription factors, such as PPRC1 and c-myc. Overall, this leads to reduced mitochondrial biogenesis resulting in a metabolic switch with concomitant anti-inflammatory effects. These data are consistent with an opposing effect of GSK-J4 on Th17 T-cell differentiation pathways directly related to proliferation and effector cytokine profiles.

**SIGNIFICANCE STATEMENT:** The precise regulation of Th17 cell metabolic function is central to an inflammatory response. Following activation, T cells undergo metabolic reprogramming and utilise up-regulated glycolysis to increase the availability of ATP. This work establishes an epigenetic link between the H3K27 demethylases KDM6A/B and the coordination of a metabolic response in activated Th17 cells. Inhibition of KDM6A/B leads to global increases in the repressive H3K27me3 histone mark resulting in down-regulation of key transcription factors, followed by metabolic reprogramming and the induction of anergy in Th17 cells. This work suggests a critical role of KDM6 enzymes in maintaining Th17 functions by controlling metabolic switches, necessary for T cells to adapt to their specific roles.

## INTRODUCTION

Upon activation, naïve CD4^+^ T cells undergo differentiation into different types of T helper cells, including T helper 1 (Th1), Th2 and Th17, characterised by different cytokine profiles and effector functions (1). In addition to their role as potent inducers of tissue inflammation, Th17 cells also serve physiological roles in host defence against bacterial or fungal infections and provide key roles in maintaining tissue barrier homeostasis (2-5). Transforming growth factor beta (TGF-β) and interleukin 6 (IL-6) are cytokines pivotal in the initial commitment of naïve T-cells towards the Th17 lineage (6-8), supported by other cytokines such as IL-23, IL-21 and IL-1β, involved in the expansion, stability and function of Th17 cells (9-11). The master transcriptional regulator RAR-related orphan receptor gamma (RORγt) is necessary to coordinate the Th17 commitment and promotes IL-17 expression (12). A loss or inhibition of RORγt can block Th17 differentiation *in vitro* and is inhibitory in models of autoimmune encephalomyelitis, imiquimod-induced cutaneous inflammation and collagen-induced arthritis (13-15).

The critical role of epigenome and chromatin modifications in differentiation and commitment from naïve T cells to T helper subsets has been demonstrated (16, 17). Chromatin templated post-translational modifications control gene transcription, DNA replication and repair, resulting in the concept of a chromatin or “histone code” that determines distinct cell states (18). In this context histone methylation plays a fundamental role in the maintenance of both active and suppressed states of gene expression, depending on the sites and degree of methylation (19). The *trithorax* and *polycomb* paradigms in particular, identified in Drosophila genetics, are centered around tri-methylation of histone H3 at lysine residues −4, (H3K4me3) which is implicated in the activation of transcription, and methylation of histone H3 at lysine −27 (H3K27me3), which is correlated with repression of transcription. The reversibility and dynamic behaviour of H3K27 methylation is provided by the methyltransferase *enhancer of zeste homolog 2* (EZH2) and by several members of the Jumonji domain containing (Jmj) Fe2+ and 2-oxoglutarate dependent oxygenases, which catalyse demethylation of methylated histone lysine residues *in vitro* and *in vivo*. In particular, Jmj family members 3 (JMJD3, KDM6B) and Ubiquitously transcribed Tetratricopeptide repeat gene, X chromosome, (UTX, KDM6A) were shown to be specific demethylases of H3K27me2/3.

Global analysis of histone modifications and DNA methylation in different T cell subsets has led to a better understanding of the mechanisms controlling differentiation and plasticity crucial for the function of T helper subsets (20-22). Integrated analysis of epigenomic profiles supports a linear model of memory differentiation, where epigenetic mechanisms control the activation of fate determining transcription factors (22). A limited number of studies have investigated the epigenetic mechanisms involved in regulating Th17 differentiation and function. Hypomethylation of DNA cytosine residues in Th17 specific genes, IL17A and RORC, shows a strong correlation with differentiation and the activation of effector function (23). Global mapping of H3K4me3 and H3K27me3 histone marks has revealed that chromatin modifications also contribute to the specificity and plasticity of effector Th17 cells, and provides a framework for using global epigenomic analyses to understand the complexity of T helper cell differentiation (24). Subsequently, chemical screening using inhibitors against various components of the epigenetic machinery has revealed novel epigenetic pathways that regulate Th17 effector function. These include the BET bromodomains, the CBP/p300 bromodomain and the KDM6A/KDM6B Jumonji histone demethylases, able to regulate CD4+ differentiation or Th17 function *in vitro* (25-28).

Metabolic pathways are intimately linked with epigenetics, transcriptional regulation and modulate cell fate and function (29-32). Moreover, targeting metabolic pathways with small molecules in autoimmunity may be a beneficial strategy for the treatment of Th17-mediated disease, such as Ankylosing Spondylitis (AS). For example, it has been reported that metabolic reprogramming using the small molecule aminooxy-acetic acid is sufficient to shift the differentiation of Th17 cells towards an inducible regulatory T cell (iTreg) phenotype, involving accumulation of 2-hydroxyglutarate, leading to hypomethylation of the gene locus of the key Treg transcription factor *foxp3* (33).

Here we establish a novel link between the H3K27 demethylases KDM6A and KDM6B in regulating Th17 cell metabolism. We show that KDM6A and KDM6B demethylases are key factors in regulating the Th17 pro-inflammatory phenotype, and control metabolic function and differentiation into effector cells. Inhibiting these enzymes results in a global increase in H3K27me3 with consequential metabolic reprogramming that leads to the emergence of an anergic phenotype.

## RESULTS

### Inhibitor screening identifies histone H3K27 demethylases as key regulators of pro-inflammatory effector T cell phenotypes

Using a focused library of small molecule inhibitors (**SI Appendix**) that target eight major classes of chromatin and epigenetic proteins, we measured the expression of Th cytokines following 24 hours of treatment of *in vitro* differentiated CD4+ cells. We evaluated the expression of IFN-γ, IL-4 and IL-17 as markers for Th1, Th2 and Th17 cells, respectively (**Figure 1A**). A consistent downregulation of cytokine expression following treatment with inhibitors against the JmjC-domain containing histone demethylases (KDM6A/KDM6B), bromodomains and histone deacetylases (HDACs) was observed in all effector subsets (**Figure 1A, Figure S1A**), suggesting a common epigenetic susceptibility. Given the importance of H3K27 (KDM6) demethylases in cellular development (34, 35) and inflammation (36), we focused this study towards the prototypic KDM6 inhibitor GSK-J4 (37), displaying an EC_50_ of 2μM (**Figure S1B)**, and its inactive regional isomer GSK-J5 as a control. Using two different Th17 differentiation models, in which either CD4^+^ or CD45RA^+^ cells are cultured in a cocktail of IL-6, IL-23 and TGF-β cytokines for 7 days, followed by treatment with DMSO, GSK-J4 or GSK-J5 for 48 hours, we observed a significant reduction in IL-17 and IFN-γ (**Figure 1B, Figure S2A**) by GSK-J4, with constant levels of RORC expression (**Figure S2A**). In contrast, addition of GSK-J4 during the differentiation process led to reduced cytokine production (**Figure S2B, S2C**), with a significant reduction in the expression of RORC (**Figure S2C**), indicating that this Th17 master transcription factor requires KDM6 demethylase activity during differentiation and not in the effector state.

**Figure 1:**
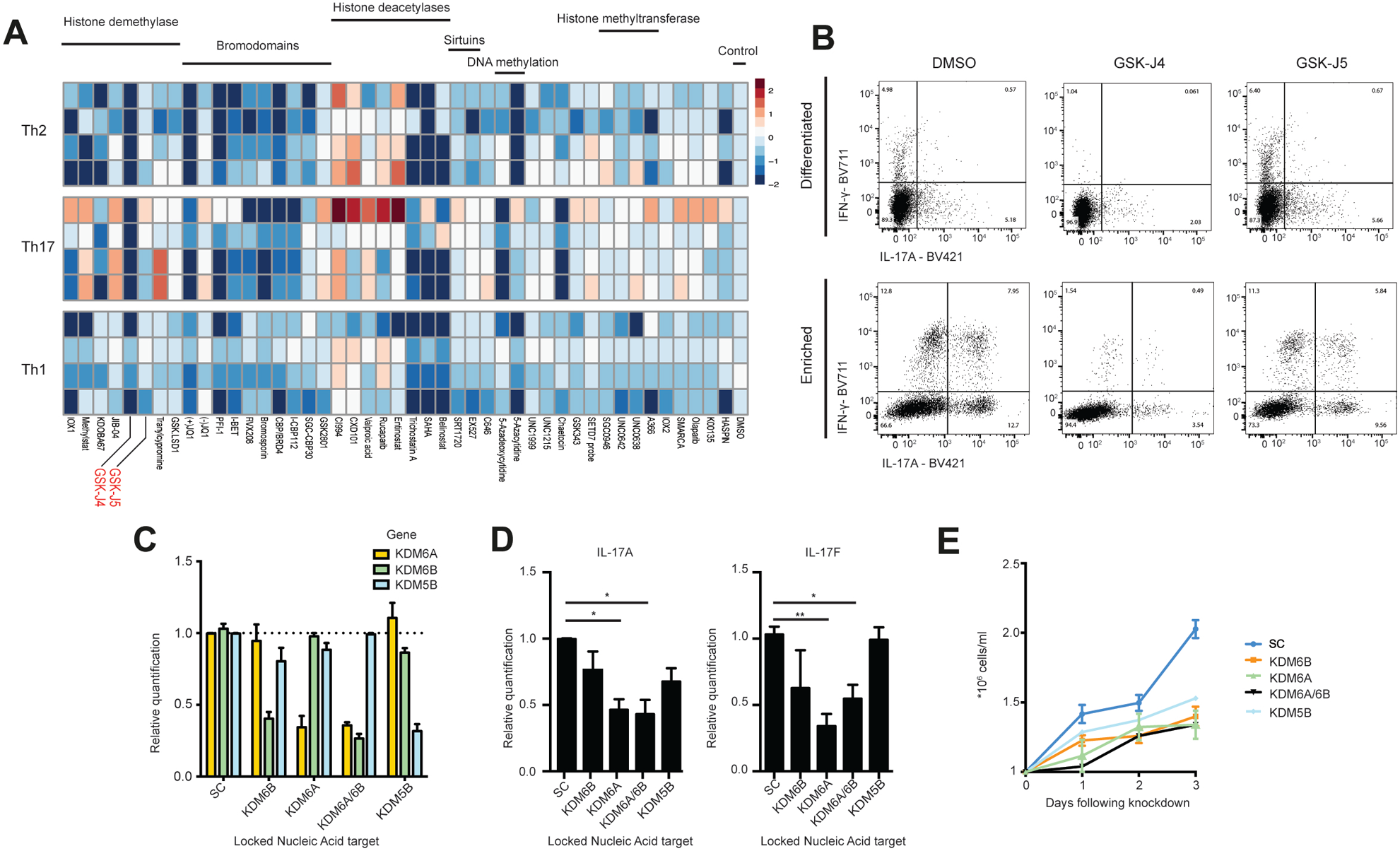
Small-molecule inhibitor screening identifies KDM6A and KSM6B as key regulators of Th17 function. (A) Heatmap reporting of Log2 fold change for secretion of IL-4, IL-17 and IFNγ from CD45RA^+^ T cells differentiated in Th2, Th17 and Th1 polarising conditions. Cytokine expression was measured by ELISA following 24 hours of culture with a library of small-molecule epigenetic inhibitors. (B) Flow cytometric analysis of IL-17 and IFN-γ staining for Th17 differentiated cells treated with DMSO, GSK-J4 or GSK-J5. (C) mRNA expression as measured by real-time PCR following 7 days of locked nucleic acid (LNA) knockdown of GSK-J4 targets, KDM6A, KDM6B and KDM5B. Data represent mean ±SEM (n=3). (D) Real time PCR quantification of IL-17A and IL-17F following LNA knockdown of KDM6A, KDM6B, dual knockdown KDM6A/KDM6B or KDM5B. (D) Cell counting following knockdown of KDM6A, KDM6B, KDM5B or a scrambled control LNA in Th17 differentiated cells over 3 days of culture. P values were calculated using a Mann-Whitney test. *P < 0.05, **P <0.01. Error bars show mean +/-SD

### Histone demethylases KDM6A and KDM6B regulate Th17 cell maturation

We observed a decrease in the activation of Th17 cells, as measured by CD25 and CCR4 flow cytometry staining, following culture in the presence of GSK-J4 (**Figure S2D and S2E)**. No increase in cell death was measured following 48 hours of GSK-J4 treatment, as measured by propidium iodide and Annexin-V staining (**Figure S2F**). GSK-J4 is a cell-permeable and potent selective inhibitor of KDM6 demethylases and has ∼5-20-fold lower activity against KDM5B demethylating enzymes in vitro (37, 38). In order to assess the specificity of GSK-J4 and evaluate the contribution of each KDM enzyme to Th17 function, we used locked nucleic acid (LNA) knockdown of KDM6A/B and the potential GSK-J4 off-target, KDM5B (JARID1B) (**Figure 1C**). Knockdown of KDM6A and KDM6B, either against single or simultaneously against both targets, led to reduced levels of IL-17 and IFN-γ (**Figure 1D, Figure S3A, S3B**). Moreover, knockdown of KDM6A and KDM6B also impacted on the proliferation of Th17 cells, as measured by cell counting (**Figure 1E**), whereas KDM5B knockdown has a lesser impact on these phenotypes. The results corroborate the role for Jumonji histone demethylase activity regulating the pro-inflammatory function and proliferation of Th17 cells (28, 39) and confirm the previously noted KDM6A/B on-target activity of the GSK-J4 chemical tool compound (37).

### KDM6 inhibitor treatment suppresses pro-inflammatory cytokine production and proliferation in autoimmune patient cells

It was of interest to validate the earlier anti-inflammatory observations within a context of chronic inflammation. Consistent with our earlier work on chronic inflammation in NK cells (36), we observe IFN-γ reduction in GSK-J4 treated CD3^+^ T cells isolated from Rheumatoid Arthritis (RA) patients (**Figure 2A**), and also of IL-17 in AS patients (**Figure 2B**). A significant reduction in the proportion of cells in G2/M phase (**Figure 2C**), associated with reduced Th17 proliferation following treatment with GSK-J4 (**Figure 2D**) was observed. The reversible nature of the GSK-J4 effect is demonstrated in washout experiments, where cells were treated for 48 hours with the inhibitor, followed by media change after 24 hours when GSK-J4 was omitted. After 48 hours cells partially regained their ability to secrete cytokines (**Figure 2E**). Single cell profiling using mass cytometry (CyTOF) allowed deep immune phenotyping and analysis of cell cycle progression of Th cells enriched from AS patients in response to GSK-J4 treatment. t-distributed stochastic neighbour embedding (t-SNE) plots (**Figure S4A**) confirm that GSK-J4 has a broad anti-inflammatory effect (36, 37, 40), highlighting a reduction in TNFα, IL-22 and IL-17 cytokines (**Figure S4B**). Activation markers such as HLA-DR and CD25 were found to be reduced following treatment with GSK-J4, in addition to a reduction in cell proliferation marker Ki-67 (**Figure S4C**). Interestingly, we observed an upregulation of CXCR3, CCR7 and CD3 surface marker expression following GSK-J4 treatment, suggesting an increase in activated and/or memory T cells (**Figure S4C**) To identify cell cycle clusters, we manually gated based on the expression of phospho-Histone H3 (P-Histone, Ser10), IDU and pRb, representing markers of cell cycle progression (**Figure 2F and Figure 2G**). This confirms a reduction in S phase and an increase in cells in G0 phase. Combined, these data suggest that GSK-J4 induces a reversible anergic state in human Th17 cells.

**Figure 2:**
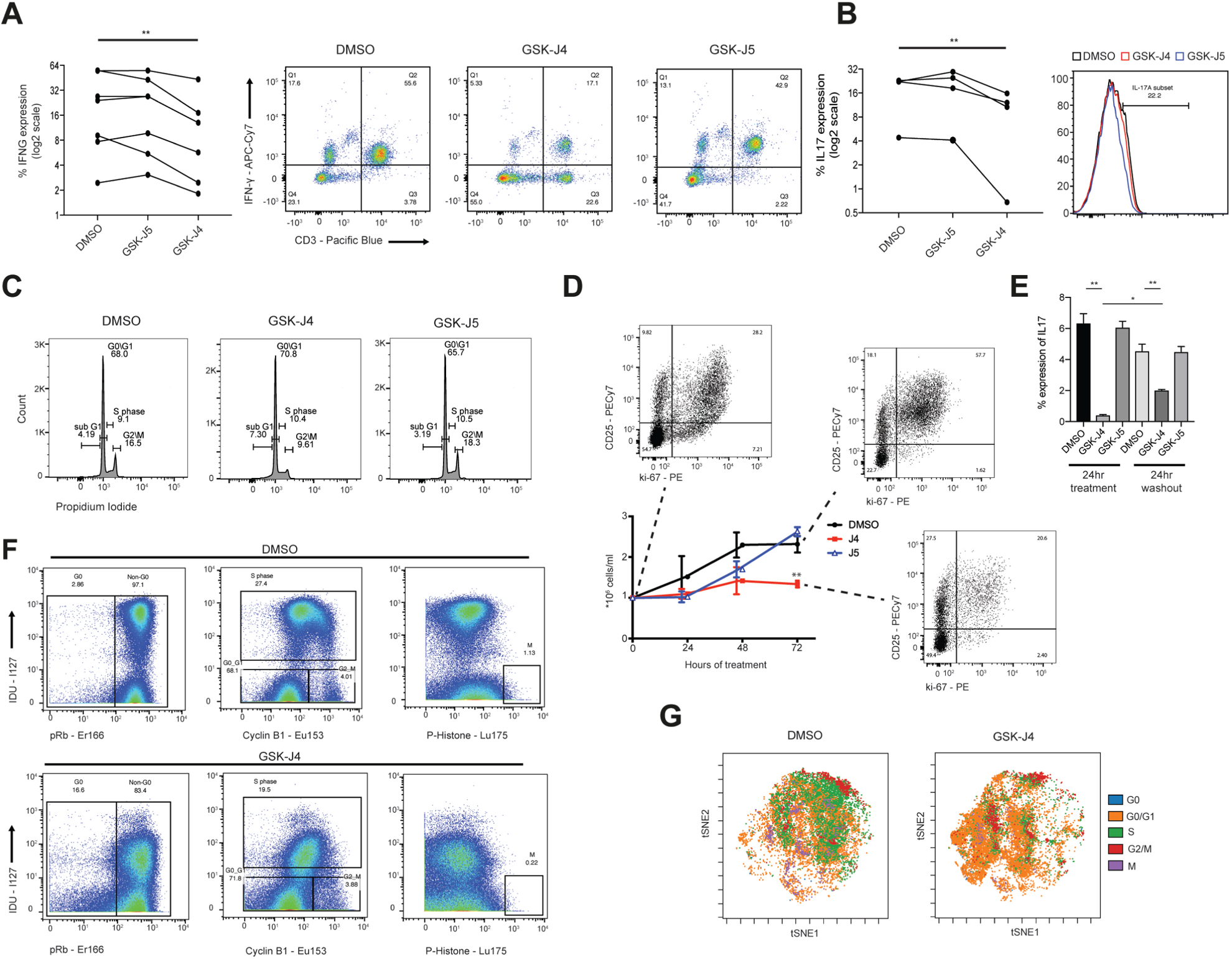
KDM6A and KDM6B regulate Th17 pro-inflammatory function in both acute and chronic inflammatory disease. (A) Flow cytometric evaluation of IFNγ expression in CD3^+^ T cells isolated from Rheumatoid Arthritis patients. Cells were treated with DMSO, GSK-J4 or GSK-J5 and stimulated for 4 hours with PMA/ionomycin in the presence of protein transport inhibitor for the last 3 hours of culture (n=7). Left panel shows percent expression of IFNγ. Right panel shows representative flow cytometry plots. (B) Flow cytometric evaluation of IL-17 expression in CD4^+^ T cells isolated from Ankylosing spondylitis patients. (C) Cell cycle evaluation using PI staining following 24 hours of DMSO, GSK-J4 or GSK-J5 treatment. The numbers in each gate represent the percent of cells in each phase of cell cycle. (D) Cell counting following 72 hours of culture with DMSO, GSK-J4 or GSK-J5. Flow cytometric analysis shows the expression of ki-67 and CD25. (E) Th17 differentiated cells were treated with DMSO, GSK-J4 or GSK-J5 for 24 hours then cells were washed three times and cultured for a further 24 hours. Expression of IL17 was measured by flow cytometry. (F) Representative example of cell cycle markers using CyTOF following 24 hours of treatment with GSK-J4 or DMSO. (G) tSNE clustering showing cell cycle stages identified by the gating strategy shown in F. All flow cytometry and CyTOF experiments were representative of n=3 independent experiments. P values were calculated using Wilcoxon matched pairs test. **P < 0.01. Error bars show mean +/-SD

### Histone demethylase treatment induces transcriptional changes affecting immune phenotype and metabolism of TH17 cells

To understand the GSK-J4 mediated phenotypic changes we initially analysed gene expression using bulk RNA sequencing (RNA-seq), performed in CD4^+^ T cells enriched for 7 days in IL-6, IL-23, and TGF-β, then cultured in the presence of GSK-J4 or DMSO for 24 hours. These data reveal a transcriptional signature that comprises >2200 genes with a significant log2-fold change and with ∼58% showing down-regulation (**Figure 3A**). Reactome pathway analysis of differentially regulated genes shows that GSK-J4 impacts on processes comprising metabolic pathways, cell cycle, respiratory chain electron transport, interleukin and cytokine signalling (**Figure 3B** and **Figure S5A**) and in addition shows upregulation of ATF4 and DDIT3, indicating a possible ATF4 mediated stress response. Although ATF4 and DDIT3 (CHOP) are upregulated, confirmed by qPCR (**Figure 3D**), typical downstream metabolic targets including e.g. argininosuccinate synthase (ASS1) or asparagine synthase (ASNS) (**Figure 3C**) are not significantly affected. Knockdown of KDM6B but not KDM6A or KDM5B revealed a similar ATF4 transcriptional response (**Figure 3E**) as GSK-J4, confirming that this effect is specifically driven by inhibition of KDM6B histone demethylase activity. Downstream ATF4/DDIT3 effects do not appear to be regulated through the canonical stress-induced eIF2a phosphorylation pathway, since ISRIB (41), a generic inhibitor of the ATF4 mediated integrated stress response (ISR) did not result in cytokine levels similar to control conditions (**Figure 3F**) and possibly explains the lack of induction of classical stress-induced metabolic ATF4 targets.

**Figure 3:**
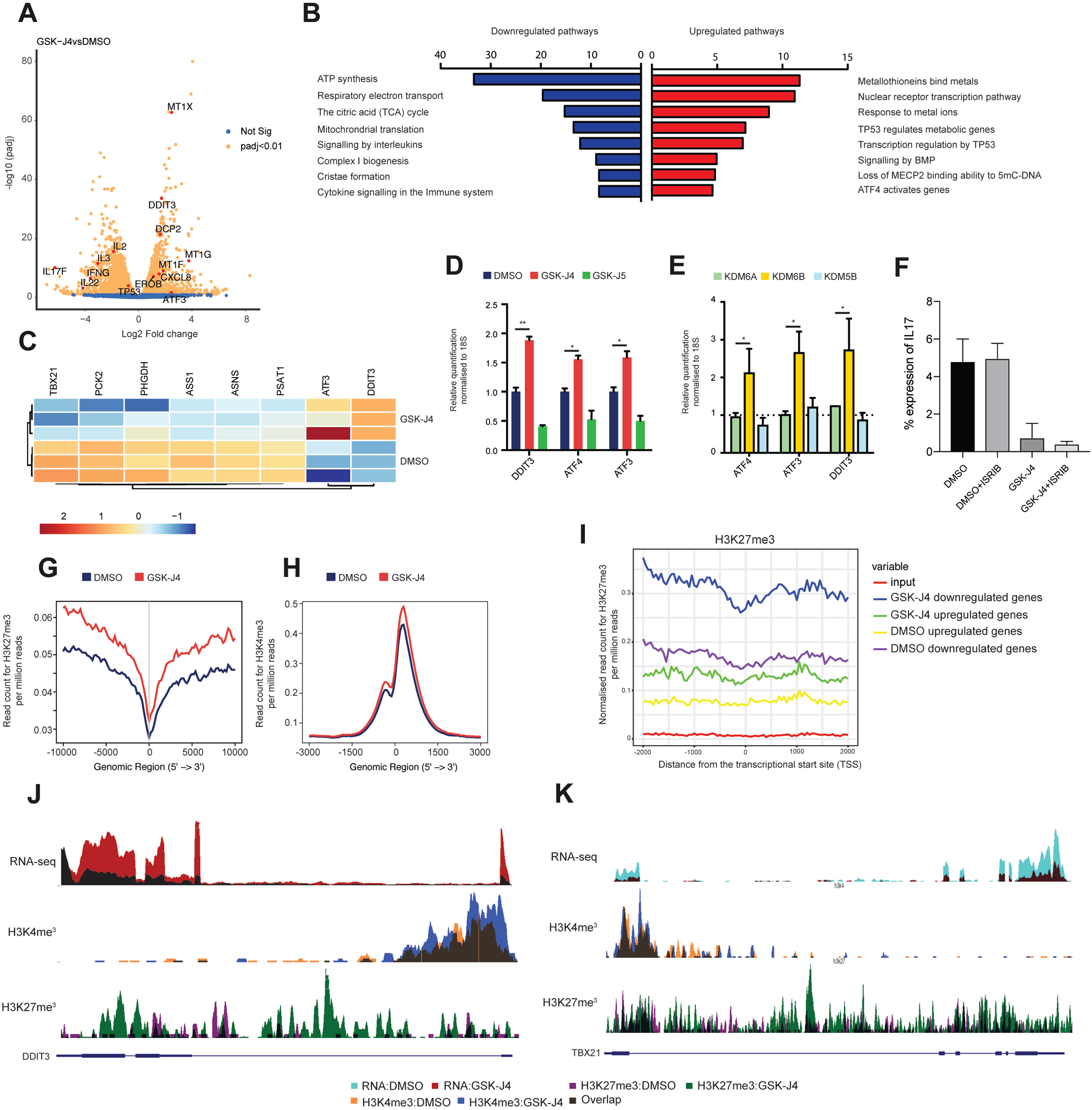
KDM6A/KDM6B inhibition induces a transcriptional change that affects immune metabolic function. A) Volcano plot showing selected gene differential expression. (B) GO analysis showing the top downregulated and upregulated ontology enrichment analysis. (C) Heatmap displaying the transcriptional response of genes associated with metabolic function. (D) RT-qPCR analysis of Th17 differentiated cells treated for 24 hours with DMSO, GSK-J4 or GSK-J5. (F) RT-qPCR analysis of Th17 differentiated cells treated for 7 days with LNAs targeting KDM6A, KDM6B or KDM5B. (G) Coverage plot showing the H3K27me3 read count per million collapsed around the TSS of all genes following treatment with DMSO or GSK-J4. (H) Coverage plot showing the H3K4me3 read count per million collapsed around the TSS of all genes following treatment with DMSO or GSK-J4. (I) Coverage plot showing H3K27me3 read count per million across upregulated and downregulated genes from Th17 differentiated cells treated with either DMSO or GSK-J4. (J) Genome browser view of the DDIT3 gene showing RNA-seq and the enrichment of H3K4me3 and H3K27me3 in cells treated with either DMSO or GSK-J4. (K) Genome browser view similar to J for TBX21 gene. P values were calculated for D and E using a Mann-Whitney test. Error bars show mean +/-SD. *p < 0.05, **p < 0.01

Next, we examined the global levels of H3K27me3, the KDM6 demethylase substrate, in addition to H3K4me3. We identified substantial global increases in H3K27me3 at transcriptional start sites (TSS) upon GSK-J4 inhibition, consistent with the expected effect of KDM6A/KDM6B as H3K27 demethylases (**Figure 3G**). In line with a large fraction of upregulated genes we also observed global increases in H3K4me3 at TSSs following GSK-J4 treatment (**Figure 3H**). The change in H3K27me3 was greater in genes that showed the largest downregulation (**Figure 3I**) supporting a model of PRC mediated gene silencing. However, several upregulated genes such as DDIT3 show inverse patterns such as increased levels of repressive H3K27 marks with reduced H3K4me3 levels (**Figure 3J**), whereas the T-box transcription factor Tbet (TBX21) regulating Tcell functions and being a direct, repressed target of DDIT3 (42), follows a pattern of PRC-mediated silencing with increased H3K27me3 levels and reduced H3K4me3 and transcript levels **(Figure 3K**).

### Single-cell transcriptomics reveals distinct populations of inflammatory Tcells following KDM6 inhibition

The observation of extensive heterogeneity with respect to cell cycle progression (**Figure 2**), prompted us to evaluate metabolic or immune gene expression by using single-cell transcriptomics. We initially performed single-cell analysis on PBMC samples cultured in the presence of either DMSO or GSK-J4 and observed a general anti-inflammatory response in all major cell types exposed to GSK-J4 (**Figure S6A**). CD4^+^ T cells were then isolated from AS patients and enriched for 7 days in IL-6, IL-23 and TGF-β, followed by 24 hours treatment with DMSO or GSK-J4. Single-cell transcriptome analysis was then performed on ∼2,000 cells from each donor (n=3 donors) (**Figure 4A**). We identified 9 unique clusters based on their gene expression profiles, comparable across all three patients (**Figure 4B, Figure S6B and S6C**). Similar to the mass cytometry data, we observed a significant shift in the t-SNE clustering following GSK-J4 treatment, with reduced numbers of cells within clusters 1, 2, 4, 8, 9 and increased numbers of cells within clusters 3, 5, 6 and 7 (**Figure 4B**). Clusters 1 and 3 showed the strongest expression of inflammatory cytokines, such as IFN-γ and IL-17A (**Figure 4C and Figure S6D**), and displayed a strong shift in cell numbers following treatment with GSK-J4. In agreement with our bulk RNA-seq data, we also observed a significant increase in DDIT3 expression following GSK-J4 treatment (**Figure 4D**). We identified discriminatory markers that account for the shift in T cell clustering in response to GSK-J4 for each of the clusters (**Figure 4E**). Clusters 1, 4, 8 and 9 were associated with a robust inflammatory signature, with enrichment of cytokines such as IL-26, IL-17A, IL-17F and IFN-γ (**Figure 4C, Figure 4E, Figure S6D**). In contrast, clusters 5 and 7, which are nearly absent in the DMSO treatment, are primarily associated with an increase in cell numbers following GSK-J4 treatment, and show a metallothionein gene enrichment and an ATF4 response (**Figure S6E and S6F**). Clusters 8 and 9 produce high levels of chemokines, with cluster 8 expressing high levels of proliferative markers such as MKI67 (Ki-67) (**Figure S6G**). Cluster 7, which is almost exclusively expressed following GSK-J4 treatment, shows high expression of T cell memory markers, such as PTPRC (CD45) and CCR7, indicating a shift from proliferating or inflammatory cells towards a memory or resting phenotype (**Figure 4E and Figure S6H)**. To further investigate the complex pattern of T cell differentiation in response to GSK-J4, we performed pseudo-temporal ordering to identify genes that are differentially regulated as a component of T cell differentiation. By using a semi-supervised method, where genes associated with the GO term T cell activation (GO:0042110) we defined the biological trajectory (43). DMSO treated and GSK-J4 treated cells are placed at opposite ends of the pseudotime trajectory (**Figure S7A**), confirming that GSK-J4 treatment leads to a significant reduction in genes associated with T-cell activation. This approach also identified a significant number of genes associated with the TCA cycle and the electron transport chain that were downregulated as a consequence of GSK-J4 treatment (**Figure S7B, supplementary data).**

**Figure 4:**
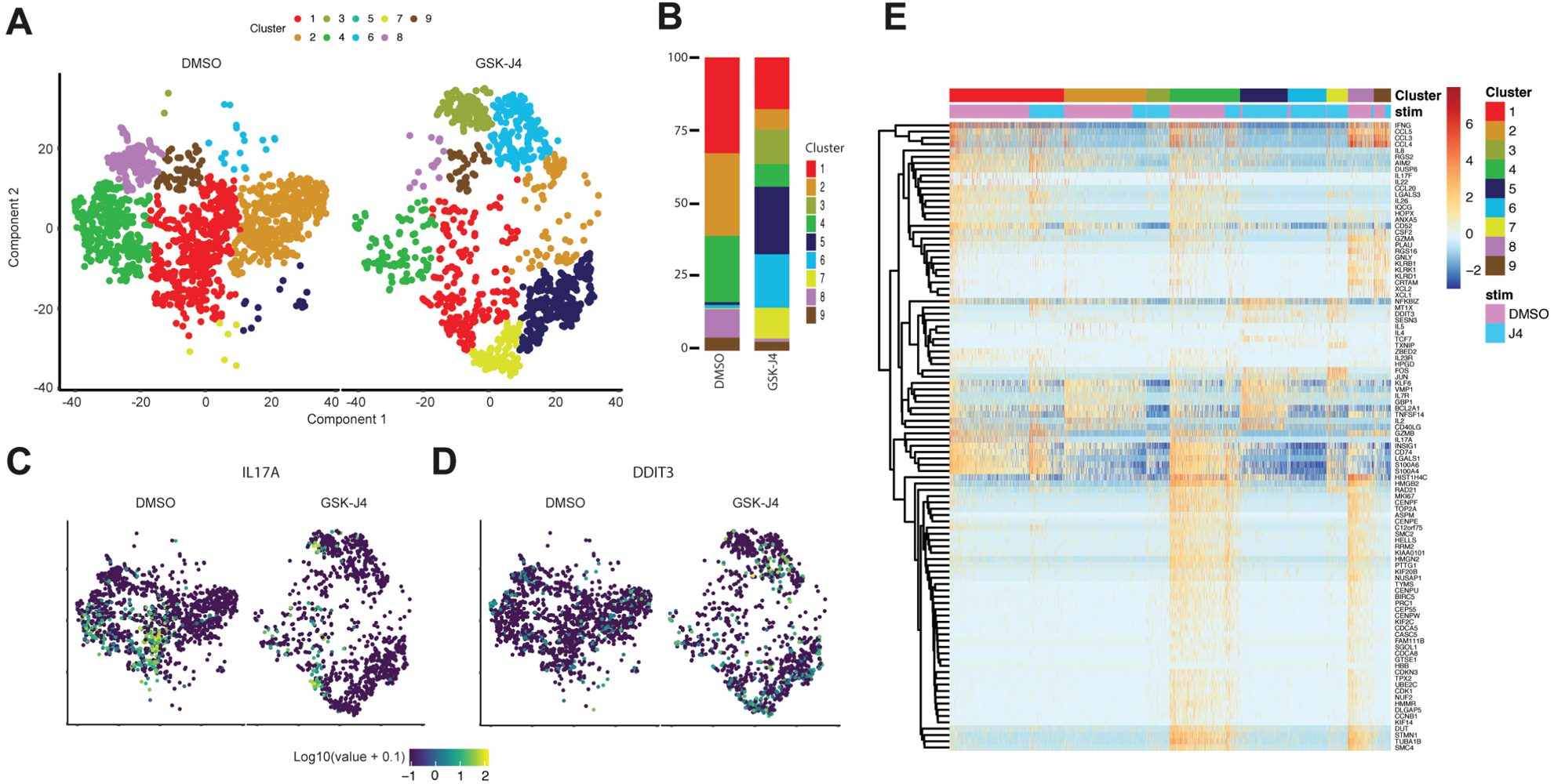
Single cell transcriptomics identifies a complex T cell response to KDM6A/KDM6B inhibition. (A) tSNE plot showing clustering of CD4^+^ T cells isolated from AS patients cultured in Th17 promoting cytokines. Left panel shows cells cultured for 24 hours in DMSO and right panel shows the cells following GSK-J4 treatment. (B) Plot showing the proportion of each cluster in both treatment groups. (C) tSNE plot showing the expression of IL-17A expression. (D) tSNE plot showing expression of DDIT3. (E) Heatmap showing the cluster specific markers for each population.

### GSK-J4 induces metabolic reprogramming in Th17 cells

the observed transcriptomic changes suggested a link between H3K27 demethylase inhibitor treatment and metabolic perturbations, leading us to investigate in more detail the resulting T cell metabolome. Human CD45RA+ T-cells were isolated from 6 healthy donors, Th17 cells were obtained as described above and treated with DMSO, GSK-J4 and its negative control GSK-J5. Extracts taken at 12 hr and 24 hr timepoints were subjected to an LC/MS based approach. Random Forest comparisons produced predictive accuracies of 83.3% or greater (DMSO vs. GSK-J4 vs. GSK-J5) (**Figure S8A**), which is well above what would be expected from random chance alone (33% for a 3-way comparison). Cellular metabolite changes during the Th17 differentiation process indicated a notable shift towards glucose utilization, increased pentose phosphate pathway metabolism (PPP) and a marked increase in nucleotides, NAD+, FAD and ADP-ribose levels (**Figure 5A and Figure S8 and Figure S9**). Depletion of several amino acids, cellular ascorbate and a strong signature of membrane lipid biosynthesis, as well as an increase in protein glycosylation pathway activity (**Figure S9** and analysis in SI Material) were also observed. These cellular profile changes are consistent with previously noted concomitant increases in both Th17 cellular proliferation and differentiation towards effector cytokine production (27). Treatment with GSK-J4, but not DMSO or GSK-J5, induced significant changes in metabolic pathways. Elevations in 3-phosphoglycerate and phosphoenolpyruvate (**Figure 5B, Figure S8**), shows that GSK-J4 increases glucose utilization at 24hrs, whereas ribose-5-phosphate and ribulose/xylulose 5-phosphate intermediates of the pentosephosphate pathway (PPP), shown to be activated with Th17 differentiation (44), were decreased following GSK-J4 treatment (**Figure 5B, Figure S8**). Decreased PPP activity would be anticipated to have multiple biological impacts on Th17 cells, including diminished NADPH production and redox status or decreased nucleotide biosynthesis, as evidenced by significant reduced levels in multiple nucleotide metabolites **(Figure S9B)**. A significant decrease in several fatty acid (propionylcarnitine, carnitine, deoxycarnitine) or branched chain amino acid metabolism intermediates was noted for GSK-J4 treatment at 12 and 24hrs (**Figure 5B, Figure S9C, SI Material**). The metabolite changes agree with the observed transcriptional changes, e.g. increased expression of glucose transporters and glycolytic key enzymes such as phosphofructokinase/bisphosphatase or lactate dehydrogenase **(Figure 5C)** explaining the increase in glycolytic activity. Conversely, downregulation of key PPP enzymes such as glucose-6-phosphate dehydrogenase (G6PD) or phosphogluconate dehydrogenase (PGD), explain the decreased flux through PPP whereas increased levels of pyruvate dehydrogenase kinase (PDK1) suggest reduction of glycolytic intermediates into the TCA cycle. Indeed, a uniquely observed effect with GSK-J4 treatment at both time points was a significant decrease in the TCA cycle metabolites α-ketoglutarate (α-KG), succinate, fumarate and malate with increased levels of glutamine and reduced glutamate (**Figure 5B, 5C, 5D, 5E, 5F)**. This suggested that alterations in α-KG-dehydrogenase or changes in glutaminolysis that feed into the TCA cycle at the level of α-KG are related to the inhibitor-mediated decrease in α-KG. However, inspection of metabolite levels reveals that GSK-J4 reduces those TCA metabolites to a similar level as seen before differentiation (**Figure S8B**). Furthermore, no transcriptional changes in α-ketoglutarate dehydrogenase (OGDH), glutamate dehydrogenase (GLUD1) or glutaminase (GLS) are observed (**Figure 5C**). This suggests that the reduction in TCA cycle metabolites reflects a general decrease in mitochondrial function and biogenesis. Indeed, this hypothesis is supported by reduced transcription of genes involved in electron transport, TCA cycle, cristae formation, protein import, mitochondrial tRNA metabolism or protein translation (**Figure 5C**), the latter also corroborated by reduction in levels of N-formylmethionine, a mitochondrial protein building block (**Figure 5B**). Reduced mitochondrial activity is also observed by tetramethylrhodamine-ethyl ester (TMRE) staining, a measure of mitochondrial membrane potential (**Figure 5G**), and a concomitant reduction of reactive oxygen species (ROS) levels following GSK-J4 treatment (**Figure 5H**). These wide-ranging mitochondrial impairments are likely related to silencing of key transcriptional regulators of mitochondrial and nuclear encoded mitochondrial genes including c-MYC, PPARGC1a and PPRC1 (**Figure 5I and Figure 5J**). Importantly, we do not observe a transcriptional response of marker genes that would indicate a mitochondrial unfolded protein response (UPR^mt^) **(Figure 5K)** (45).

**Figure 5:**
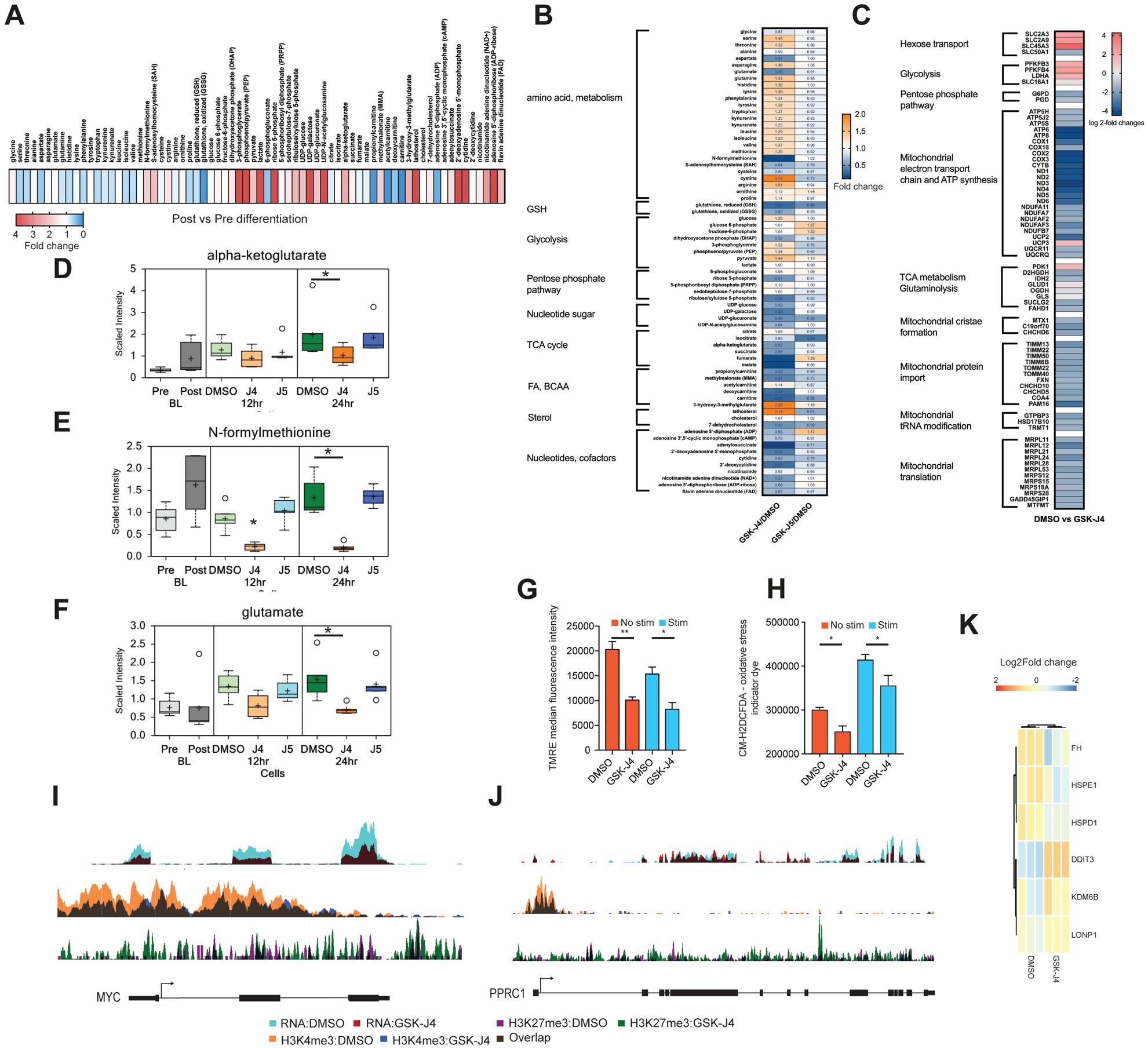
Inhibition of JMJD3/UTX leads to metabolic reprogramming of Th17 cells. (A) Heatmap showing the metabolite changes between pre-differentiation and post-differentiation. (B) Heatmap showing the expression of selected metabolites grouped into functional classes in Th17 differentiated cells treated either DMSO and GSK-J4 for 24 hours. (C) Heatmap displaying the transcriptional response of genes associated with mitochondrial metabolic regulation and function. Box plots showing the scaled intensity of (D) malate, (E) glutamine and (F) α-ketogluterate from Th17 differentiated cells treated with DMSO, GSK-J4 and GSK-J5 for 24 hours. (G) Mitochondrial membrane potential and (H) oxidative stress marker measurements following DMSO or GSK-J4 treatment of Th17 differentiated cells. (I) Genome browser view of the MYC gene showing RNA-seq and the enrichment of H3K4me3 and H3K27me3 in cells treated with either DMSO or GSK-J4. (J) Genome browser view similar to F for PPRC1 gene. (K) Heatmap showing the expression of genes involved in the UPR^mt^ response pathway. P values were calculated using a Mann-Whitney test. *p < 0.05, **p < 0.01. Error bars show mean +/-SEM.

## DISCUSSION

This work provides several new insights into epigenetic control of Th17 cell functions. Consistent with previous studies highlighting the importance of histone demethylases KDM6A/B in cellular development and disease (46-49), we here extend previous observations that these enzymes are critical regulators of Th17 development and pro-inflammatory phenotypes. In mouse CD4^+^ T cells, KDM6B ablation promotes CD4^+^ T cell differentiation into Th2 and Th17 subsets in small intestine and colon (27, 50), however two other independent studies did not observe this enhanced Th17 differentiation in KDM6B ablated CD4^+^ T cells (27, 28, 39) but showed significant impairment of Th17 differentiation and effector function. One possible explanation for this discrepancy may be related to redundant and compensatory roles of KDM6A and KDM6B in T-cell differentiation (51). In this context, our study provides clear evidence that both enzymes in humans are important regulatory elements of Th17 cells, by involvement in differentiation of Th17 cells, and in addition by controlling proliferation and pro-inflammatory effector functions of mature Th17 cells.

Single-cell transcriptomics has provided detailed pictures of cellular heterogeneity related to disease pathology (52), in which Th17 cells exhibits a range of phenotypes from pathogenic to regulatory in nature (53, 54). Our single-cell transcriptomics data indeed show significant T-cell heterogeneity, with a consistent set of phenotypes that correlate with subsets of highly inflammatory, memory and resting cells. In agreement with our bulk RNA-seq analysis, KDM6 inhibitor treatment of Th17 cells leads to a significant downregulation in Th17 specific cytokine function and to anti-proliferative effects. This is clearly reflected in the single cell data where significant shifts of highly inflammatory cells following GSK-J4 treatment are observed towards a resting state, suggesting a preferential inhibitor activity on proliferating and inflammatory cells.

Importantly, an effective Th17 cell response requires a T cell to adapt its metabolic state in response to various microenvironments and stimuli. Quiescent T-cell energy demands are low, they primarily oxidise glucose-derived pyruvate in their mitochondria, which ensures optimal ATP production per molecule of glucose (55, 56). Following activation, T cells undergo metabolic reprogramming in order to significantly increase the availability of ATP. These are usually short-lived events that rely primarily on glycolysis for their energy demands (55, 56), which is also observed in cytokine-activated lymphocytes (57). Supporting our observations, an elevated rate of aerobic glycolysis is accompanied by increased expression of glycolytic enzymes and nutrient transporters (58). This increased flux through the glycolytic pathway is controlled by mTORC1 signalling and provides the necessary metabolic precursors for proliferation and effector functions (59).

A conspicuous transcriptional response to GSK-J4 treatment, corroborated by knockdown experiments, is a change in metabolic gene expression. Although we observe a characteristic ATF4 signature in NK cells following GSK-J4 treatment (36), we only observe a robust DDIT3 upregulation in this study with a consistent lack of upregulated metabolic targets, which are otherwise associated with an ATF4-DDIT3 mediated stress response. Importantly, DDIT3 has been shown to directly repress critical Tcell transcription factors like T-bet, leading to reduced cytokine output in tumour-infiltrating Tcells and thus might contribute to the observed anti-inflammatory phenotype (42).

The observed metabolic changes-i.e. upregulated glycolytic and downregulated TCA cycle activity leading to consequent reduction in inflammatory cytokine production and anergy- are best explained by the observed H3K27 mediated silencing of key transcription factors such as MYC, PPARg and PPRC1 as a consequence of GSK-J4 treatment (61). Recently, it has been shown that inhibition of mTORC1 signalling rescues ATF4-deficient cells from MYC-induced endoplasmic reticulum stress (60). This reveals an essential role for ATF4 in survival following MYC activation and our data suggests that KDM demethylases are an important regulator of this pathway.

We postulate that these mitochondria-centered events, i.e. reduced biogenesis and function, and the TCR stimulated increase in glycolytic activity are critical drivers of the observed Th17 phenotype upon demethylase inhibition where anergy is associated with the development of a memory T cell phenotype that leads to downregulation of glutaminolysis and lipid biosynthesis (62). Importantly, the observed mitochondrial impairment upon demethylase inhibition does not show any sign of stress such as the mitochondrial unfolded protein response (UPR^mt^), often associated with a dysfunctional electron transport chain, increased ROS production and an altered transcription factor as well as a distinct effector profile (63, 64). Although previous work in C.elegans has demonstrated a critical involvement of histone demethylases, including the KDM6B ortholog JMJD3.1, and controlling UPR^mt^ effector regulation we here do not correlate KDM6B function with a mitochondrial stress response such as UPR^mt^ (65).

Taken together, our results suggest a critical role of KDM6 enzymes in maintaining Th17 functions by controlling metabolic switches, necessary for Tcells to adapt to their specific roles. The results provide a basis for further investigations into using small molecule epigenetic and metabolic inhibitors to understand these critical facets of the adaptive immune system.

## MATERIALS and METHODS

### Lymphocyte isolation

T cells were isolated from venous blood, obtained from healthy volunteers, or from single-donor platelet pheresis residues from the Oxford National Health Service Blood Transfusion Service. Peripheral blood was obtained from RA patients attending the rheumatology clinic at Northwick Park Hospital, London. Peripheral blood obtained from AS patients was collected at the Nuffield Orthopaedic Centre Hospital, Oxford. The study was approved under the Research Ethics Committee (REC) Number 07/H0706/81 and Oxford Research Ethics committee (number 06/Q1606/139). The human biological samples were sourced ethically, and their research use was in accord with the terms of the informed consents. The patients were diagnosed according to the American College of Rheumatology (ACR) Eular 2010 criteria. Mononuclear cells were isolated by Ficoll density gradient centrifugation and CD4^+^ or CD45RA^+^ T cells were isolated using magnetic bead isolation using Dynabeads (Invitrogen).

### T helper cell differentiation and enrichment

Human CD4^+^ T cell subsets *in vitro* differentiation was performed as previously described (66-68). Briefly, naïve CD4^+^ CD45RA^+^ T cells were purified from PBMCs of healthy donors with T-cell enrichment kit (Miltenyi) and then cultured in the presence of anti-CD3/CD28 activation beads (1:20 bead to cell ratio) (Invitrogen) plus polarization condition medium of RPMI-1640 containing 10% knockout serum replacement (ThermoFisher) supplemented with the following cytokines at 37C for 6 days. For Th1 differentiation, naïve T cells were cultured in the presence of anti-IL-4 neutralizing antibody (10 mg/ml, R&D), and recombinant human IL-12 (5 ng/ml, R & D). For Th2 differentiation, naïve T cells were cultured in the presence of anti-IFN-g neutralizing antibody (10 mg/ml, R&D) and recombinant IL-4 (4 ng/ml, Peprotech). For Th17 differentiation, naïve T cells were cultured in the presence of IL-1b (20 ng/ml, Peprotech), IL-6 (20 ng/ml, Peprotech), recombinant TGF-b (3 ng/ml, Peprotech) and IL-23 (10 ng/ml) (R & D) for 7 days. For Th17 cell expansion, CD4^+^ T cells were isolated using Dynabeads™ (Invitrogen) as opposed to CD45RA^+^ T cells.

### RT-qPCR Analysis

Total RNA was extracted from T cells using Trizol reagent (Invitrogen), and cDNA was transcribed using a SuperScript II RT kit (Invitrogen), both according to manufacturers’ instructions. Expression levels of each gene were determined by reverse-transcription PCR using specific primers, and mRNA levels in each sample were normalized to the relative quantity of b-actin gene expression. All experiments were performed in triplicate. The specific primers used in this study are listed in Table S3.

### Compound inhibitor screens

Following Th cell differentiation, the cells were seeded at 50,000 cells/well in 200 μL of culture medium in 96 well plates. Compounds were dissolved in DMSO at concentrations previously described in Cribbs *et al* 2018 and diluted to achieve the desired working solutions. Compound effects were compared with cells cultured in 0.1% DMSO alone, whereas wells filled with media served as a background control. Following 24 hours of compound treatment, media was harvested and Enzyme-linked Immunosorbent assay (ELISA) was used to measure cytokine release. Small molecules that were able to modify cytokine expression by a 2-fold change and P < 0.05 were considered as significant.

### Locked nucleic acid knockdown experiments

Knockdown experiments were conducted using locked nucleic acids (LNA) antisense oligonucleotides designed and synthesized as described previously (69) by Santaris A/S. Following synthesis oligonucleotides were HPLC purified, desalted using a Milliprep membrane, and verified by LC-MS. Th cells were cultured at a concentration of 1 x 106 cells/ml, stimulated with IL-15, and LNAs were added to the culture media at a concentration of 1 μM, then cultured for a further 8 days.

### Enzyme-linked Immunosorbent assay (ELISA)

IFN-γ, IL17 and IL-4 secretion were determined in the cell culture supernatant by ELISA (eBioscience). The assay was performed according to the manufacturer’s instructions, and all standards and samples were measured in triplicate.

### Chromatin immunoprecipitation followed by sequencing

Per chromatin immunoprecipitation experiment, 30 × 10^6^ total T cells were used. All ChIP-seq assays were performed using single donors. For quantitative ChIP experiments, 4 × 10^6^ SF1 cells were spiked into the pool. The cells were then cross-linked by formaldehyde treatment, and chromatin was fragmented to 200–300 bp by sonication using a Biorupter® Pico (Diagenode). Each lysate was immunoprecipitated with 10 μg of the following antibodies: H3K4me3 (Merck Millipore) and H3K27me3 (Merck Millipore). The ChIP was then performed for each antibody as described previously by Orlando *et al* (70). Purified DNA was used for library preparation using a NEBNext Ultra DNA sample preparation kit (NEB) according to the manufacturer’s recommendations. The samples were multiplexed, quantified using a Kapa library quantification kit (KAPA Biosystems), and sequenced on a NextSeq 500 (Illumina) platform (paired-end, 2 × 41 bp). Sequencing depth was in excess of 20 million reads/sample to achieve sufficient coverage, which was assessed bioinformatically.

### Flow cytometry

For flow cytometry detection of surface molecules, T cells were first incubated with anti-Fc receptor Ab, to reduce non-specific binding, and then labelled with appropriate fluorescent mAbs. For intracellular staining of cytokines, T cells were stimulated with phorbol 12-myristate 13-acetate (50 ng/ml) and ionomycin (1 mg/ml) for 4 h in the presence of protein transport inhibitor solution (WHICH ONE). The cells were collected, washed, fixed, permeabilized (fix/perm, Biolegend), and stained with fluorescein-labelled cytokine-specific mAbs according to the manufacturer’s instructions. Appropriate fluorescein-conjugated, isotype-matched mAbs were used as negative controls. Flow cytometry antibodies used in this study are detailed in the key resources table. Cells were then analysed using a BD Fortessa flow cytometry instrument.

### RNA isolation and bulk RNA-seq library preparation

RNA was isolated from NK cells using the Quick-RNA MiniPrep kit (Zymo) according to the manufacturer’s protocol. The quality of the RNA samples was verified by electrophoresis on Tapestation (Agilent). The RIN scores for all samples were in the range of 7.5–9.5. RNA-seq libraries were prepared using the NEBNext® Ultra™ RNA library prep kit for Illumina® using TruSeq indexes, according to the manufacturer’s protocol. The resulting libraries were sequenced on a NextSeq 500 platform (Illumina) using a paired-end run 2 × 80 bp, to an average depth of 112 × 10^6^ paired-end reads/sample (range 47 × 10^6^ to 168 × 10^6^).

### Single-cell RNA-seq library preparation

Single-cell capture and reverse transcription (RT) were performed as previously described (71). Cells were loaded into a microfluidics cartridge at a concentration of 310 cells per uL. Cell capture, lysis and reverse transcription were all performed using a Nadia instrument (Dolomite Bio). RT reactions were performed using ChemGene beads also captured in the microfluidic wells. Beads were collected from the device and then cDNA amplification was performed. Prior to PCR, the beads were treated with Exo-I. Following PCR purified cDNA was used as an input for Nextera tagmentation reactions. cDNA library quality was assessed using a TapeStation (Agilent Technologies). All cDNA purification steps were performed using Ampure XP beads (Beckkman). High quality samples were then sequenced on a NextSeq 500 sequencer using a 75 cycle High Output kit (Illumina).

### Metabolomic studies

Naive CD4+ T-cells were isolated from 6 human donors and subdivided into 8 equal groups: Baseline (Pre-Differentiation, undifferentiated T-cells), Th17 differentiated T-cells (Post Differentiation), Th17 differentiated cells treated with vehicle (DMSO), GSK-J4 or GSK-J5 (2 μM each in 0.1% DMSO final concentration) for both 12 and 24hrs. Following each treatment T-cells were isolated, pelleted, and snap-frozen for further global, unbiased metabolomic analysis on the HD4 accurate mass platform (72). In addition, media samples were collected for Pre-Differentiation, Post-Differentiation, and 24hrs samples treated with DMSO, GSK-J4 and GSK-J5.

### Mitochondrial function

T cells were stained with 20 µM CM-H_2_DCFDA, 200 nM or 100 nM TMRE in PBS for 30 minutes at 37°C (5% CO_2_) to determine ROS production and mitochondrial membrane potential respectively. After staining, cells were washed with PBS and fluorescence was measured using a Beckman Coulter CytoFLEX flow cytometer.

### Bulk RNA sequencing analysis

A computational pipeline was written calling scripts from the CGAT toolkit (73, 74) to analyze the next generation sequencing data and (https://github.com/cgat-developers/cgat-flow). For bulk RNA-seq experiments, sequencing reads were mapped to the reference human genome sequence (GRCh37 (hg19) assembly) using hisat v0.1.6 (75). To count the reads mapped to individual genes, the program featureCounts v1.4.6 was used (76). Only uniquely mapped reads were used in the counting step. The counts table that was generated was then used for differential expression analysis. Differential gene expression analysis was performed with DESeq2 v1.12.3 within the R statistical framework v3.3.0 (77). To define differentially expressed genes, a threshold of ±2-fold change, and a false discovery rate of <0.05 were used.

### Single cell RNA sequencing analysis

Reads were demultiplexed, aligned to the GRCh37 (hg19) assembly reference genome, and filtered; and cell barcodes and UMIs were quantified using dropseq tools (71). Dropseq tools uses STAR (78) for alignment. For each gene, UMI counts of all transcript isoforms are summed to obtain a digital measure of total gene expression. All further filtering was performed with Seurat v2.1, 1998 cells (DMSO) and 1962 (GSK-J4) were identified as good quality and were processed for downstream analyses. Of these cells, 2835 median number of genes were detected, with 430 genes expressed in at least 50% of cells. Briefly, outliers were detected for each quality metric that included cells that expressed a minimum of 50 genes per cell and a mitochondrial content of less than 5%.

### Dimensionality reduction and clustering

Full descriptions on the clustering procedure can be found in the *monocle2* documentation. Briefly, after QC and filtering, gene expression for each cell was normalised by total transcript count and log transformed. To identify highly variable genes that account for cellular heterogeneity, co-varying genes were reduced using the tSNE reduction method. For unbiased clustering, monocle2 uses density peak clustering. Clusters were then projected onto a tSNE plot. Cluster identification was robust across a range of PCs and resolutions. **Pseudotime ordering -** Pseudotime ordering was performed using the *monocle* package (v2.11.3). To isolate a set of ordering genes that define the progression of T cell activation, using the GO term “T cell activation”; GO:0042110. Next, we reduced data dimensionality using the *DDRTree* method and ordered cells along the resulting trajectory. Finally, we identified genes that change as a function of the computed pseudotime using the *differentialGeneTest* function. A *q*-value threshold of 0.01 was applied to all differential expression tests in the *monocle* workflow. Gene expression heat maps were produced using the *pheatmap* package (v1.0.12).

### Differential gene expression

Differentially expressed gene analysis was performed using the nonparametric Wilcoxon test on log_2_(TPM) expression values for the comparison of expression level and Fisher′s exact test for the comparison of expressing cell frequency. *P* values generated from both tests were then combined using Fisher’s method and were adjusted using Benjamini– Hochberg (BH). Differentially expressed genes were selected on the basis of the absolute log_2_ fold change of ≥1 and the adjusted *P* value of <0.05. Selected genes were subjected to the hierarchical clustering analysis using Pearson correlation as a distance, with clustering performed in R.

### ChIP and ATAC sequencing analysis

Bowtie software v0.12.5 was used to align the reads to the human hg19 reference genome (79). Reads were only considered that were uniquely aligned to the genome with up to two mismatches. For quantitative ChIP, the number of reads mapping to SF9 cells was determined for each sample. Bedtools version 2.2.24 was used to generate bedgraph files from the mapped BAM files using the scaling factor derived from the SF9 read count. Averaged tracks for each condition were produced representing the mean of the scaled values for biological replicates. Homer tag directories were then produced using the raw read function, and coverage plots around the TSS were plotted in R. MACS software (v1.4.2) was used to identify enrichment of intervals of H3K4me3 and H3K27me3 following ChIP-seq and regions of open chromatin following ATAC-seq. Sequencing of the whole cell extract was performed to determine the background model when analysing ChIP-seq

### Metabolomic analysis

Welch’s two-sample *t*-tests, Repeated Measures ANOVA and Random Forest analysis was used to analyze the data. For all analyses, following normalization to protein for cells, missing values, if any, were imputed with the observed minimum for that particular compound. The statistical analyses were performed on natural log-transformed data (See Appendix “Metabolite Quantification and Data Normalization”). When comparing GSK-J4 or GSK-J5 treated cells with time-matched DMSO controls using student t-tests, it became clear that a donor effect was limiting in terms of understanding the true drug related changes. Hence repeated measures ANOVA was used to compare each donor response to drug with its own DMSO control, thereby addressing the donor effect and revealing the true drug response profiles within this study (SI Material 1).

### Mass cytometry analysis

For each sample, 1-3 million cells were first stained with a solution containing rhodium DNA intercalator (Fluidigm) to distinguish live/dead and IdU (Fluidigm) to distinguish cells in S-phase, prior to Fc receptor blocking (Miltenyi Biotec). Samples were then stained with a mixture of metal conjugated antibodies recognising cell surface antigens (all antibodies purchased from Fluidigm, see supplementary data for antibodies and labels). After washing in Maxpar cell staining buffer (Fluidigm), samples were fixed in ice cold methanol for 15 minutes prior to washing and incubation with metal conjugated antibodies recognising intracellular and phospho-protein antigens. Samples were washed twice in cell staining buffer, fixed by incubation with 1.6% PFA (Pierce) for 10 minutes and finally incubated overnight with iridium DNA intercalator in Maxpar fix and perm buffer (Fluidigm). Prior to acquisition samples were washed twice in Maxpar cell staining buffer and twice in Maxpar water and filtered through a 40 um cell strainer before being acquired on a Helios mass cytometer (Fluidigm).

After acquisition, all .fcs files in the experiment were normalised using tools within the Helios software and then uploaded to Cytobank (www.cytobank.org) for all gating and further analysis including using clustering and dimensionality reduction algorithms such as SPADE and viSNE (80).

### Statistical analysis

All other statistical analyses were performed with GraphPad Prism7 software. Unless indicated otherwise, data are expressed as mean ± standard deviation (SD). D’Agostino and Pearson test was used to test whether the data come from a Gaussian distribution. For multiple group comparison, the one-way analysis of variance (ANOVA) was used, followed by the Dunnett’s test for comparing experimental groups against a single control. For single comparison between two groups, paired Student’s *t* test was used. Nonparametric t-test was chosen if the sample size was too small and not fit Gaussian distribution. The statistical parameters can be found in the figure legends.

## Supporting information

Bulk RNA-seq results

Single cell cluster differential expression

Metabolomic results

## DATA AND SOFTWARE AVAILABILITY

Single-cell RNA-seq, RNA-seq, ATAC-seq, and ChIP-seq data sets are deposited with the GEO database under accession number GSE127767. R scripts used for data analysis are available upon request.

## ACKNOWLEDGMENTS

The study was supported through funding from the Kennedy Trust for Rheumatology Research, Arthritis Research UK (program grant number 20522), the NIHR Oxford Biomedical Research Unit, Cancer Research UK, and the LEAN program grant from the Leducq Foundation. MW is supported by grants from the Netherlands Heart Foundation (GENIUS2), the Netherlands Heart Foundation and Spark Holland (2015B002) and the European Union (ITN grant EPIMAC). APC was supported by the Medical Research Council (MRC) CGAT program (G1000902) and a CRUK Oxford Centre Development Fund award (CRUKDF-0318-AC(AZ)). The Structural Genomics Consortium is a registered charity (number 1097737) that receives funds from Abbvie, Bayer Healthcare, Boehringer Ingelheim, the Canadian Institutes for Health Research, the Canadian Foundation for Innovation, Eli Lilly and Company, Genome Canada, the Ontario Ministry of Economic Development and Innovation, Janssen, the Novartis Research Foundation, Pfizer, Takeda, and the Wellcome Trust. The research leading to these results has received funding from the People Programme (Marie Curie Actions) of the European Union’s Seventh Framework Programme (FP7/2007-2013) under REA grant agreement n° [609305].

## AUTHOR CONTRIBUTIONS

AC and UO conceived and supervised the study and wrote the manuscript, AC, ST, DX, carried out experiments, MP, AC carried out sequencing experiments and analysed data, ML, SO, HO designed and provided LNA reagents, BS carried out metabolomics experiments and analysed data, JB and MW performed and supervised immunometabolomic assays, RP, NA, MF, provided reagents and materials, and PW provided AS patient material and information.

## REFERENCES

1. Sandquist I & Kolls J (2018) Update on regulation and effector functions of Th17 cells. F1000Res 7:205.

2. Korn T, Bettelli E, Oukka M, & Kuchroo VK (2009) IL-17 and Th17 Cells. Annu Rev Immunol 27:485–517.

3. Dong C (2008) TH17 cells in development: an updated view of their molecular identity and genetic programming. Nat Rev Immunol 8(5):337–348.

4. Gaffen SL, Jain R, Garg AV, & Cua DJ (2014) The IL-23-IL-17 immune axis: from mechanisms to therapeutic testing. Nat Rev Immunol 14(9):585–600.

5. Akitsu A & Iwakura Y (2018) Interleukin-17-producing gamma delta T (gamma delta 17) cells in inflammatory diseases. Immunology 155(4):418–426.

6. Bettelli E, et al. (2006) Reciprocal developmental pathways for the generation of pathogenic effector TH17 and regulatory T cells. Nature 441(7090):235–238.

7. Mangan PR, et al. (2006) Transforming growth factor-beta induces development of the T(H)17 lineage. Nature 441(7090):231–234.

8. Veldhoen M, Hocking RJ, Atkins CJ, Locksley RM, & Stockinger B (2006) TGFbeta in the context of an inflammatory cytokine milieu supports de novo differentiation of IL-17-producing T cells. Immunity 24(2):179–189.

9. Korn T, et al. (2007) IL-21 initiates an alternative pathway to induce proinflammatory T(H)17 cells. Nature 448(7152):484–487.

10. Chung Y, et al. (2009) Critical regulation of early Th17 cell differentiation by interleukin-1 signaling. Immunity 30(4):576–587.

11. Cua DJ, et al. (2003) Interleukin-23 rather than interleukin-12 is the critical cytokine for autoimmune inflammation of the brain. Nature 421(6924):744–748.

12. Ivanov II, et al. (2006) The orphan nuclear receptor ROR gamma t directs the differentiation program of proinflammatory IL-17(+) T helper cells. Cell 126(6):1121–1133.

13. Al-Mossawi MH, et al. (2014) In-Vitro Supression of Th17 Responses in Inflammatory Arthritis Patients Using Small Molecule Ror-Gamma-T Inhibitors. Ann Rheum Dis 73:357–357.

14. Huh JR & Littman DR (2012) Small molecule inhibitors of ROR gamma t: Targeting Th17 cells and other applications. Eur J Immunol 42(9):2232–2237.

15. Skepner J, et al. (2014) Pharmacologic Inhibition of ROR gamma t Regulates Th17 Signature Gene Expression and Suppresses Cutaneous Inflammation In Vivo. J Immunol 192(6):2564–2575.

16. Hirahara K, et al. (2011) Helper T-cell differentiation and plasticity: insights from epigenetics. Immunology 134(3):235–245.

17. Durek P, et al. (2016) Epigenomic Profiling of Human CD4(+) T Cells Supports a Linear Differentiation Model and Highlights Molecular Regulators of Memory Development. Immunity 45(5):1148–1161.

18. Jenuwein T & Allis CD (2001) Translating the histone code. Science 293(5532):1074–1080.

19. Greer EL & Shi Y (2012) Histone methylation: a dynamic mark in health, disease and inheritance. Nat Rev Genet 13(5):343–357.

20. Ichiyama K, et al. (2015) The Methylcytosine Dioxygenase Tet2 Promotes DNA Demethylation and Activation of Cytokine Gene Expression in T Cells (vol 42, pg 613, 2015). Immunity 42(6):1214–1214.

21. Wei Y, et al. (2013) Global H3K4me3 genome mapping reveals alterations of innate immunity signaling and overexpression of JMJD3 in human myelodysplastic syndrome CD34+cells. Leukemia 27(11):2177–2186.

22. Durek P, et al. (2016) Epigenomic Profiling of Human CD4(+) T Cells Supports a Linear Differentiation Model and Highlights Molecular Regulators of Memory Development. Immunity 45(5):1148–1161.

23. Cohen CJ, et al. (2011) Human Th1 and Th17 Cells Exhibit Epigenetic Stability at Signature Cytokine and Transcription Factor Loci. J Immunol 187(11):5615–5626.

24. Wei G, et al. (2009) Global Mapping of H3K4me3 and H3K27me3 Reveals Specificity and Plasticity in Lineage Fate Determination of Differentiating CD4(+) T Cells. Immunity 30(1):155–167.

25. Hammitzsch A, et al. (2015) CBP30, a selective CBP/p300 bromodomain inhibitor, suppresses human Th17 responses. Proc Natl Acad Sci U S A 112(34):10768–10773.

26. Mele DA, et al. (2013) BET bromodomain inhibition suppresses TH17-mediated pathology. J Exp Med 210(11):2181–2190.

27. Liu Z, et al. (2015) The histone H3 lysine-27 demethylase Jmjd3 plays a critical role in specific regulation of Th17 cell differentiation. J Mol Cell Biol 7(6):505–516.

28. Li Q, et al. (2014) Critical role of histone demethylase Jmjd3 in the regulation of CD4+ T-cell differentiation. Nat Commun 5:5780.

29. Wellen KE & Thompson CB (2012) A two-way street: reciprocal regulation of metabolism and signalling. Nat Rev Mol Cell Bio 13(4):270–U271.

30. Lu C & Thompson CB (2012) Metabolic Regulation of Epigenetics. Cell Metab 16(1):9–17.

31. Kaelin WG & McKnight SL (2013) Influence of Metabolism on Epigenetics and Disease. Cell 153(1):56–69.

32. Nowak RP, et al. (2016) Advances and challenges in understanding histone demethylase biology. Curr Opin Chem Biol 33:151–159.

33. Xu T, et al. (2017) Metabolic control of T(H)17 and induced T-reg cell balance by an epigenetic mechanism. Nature 548(7666):228-+.

34. Shpargel KB, Starmer J, Yee D, Pohlers M, & Magnuson T (2014) KDM6 demethylase independent loss of histone H3 lysine 27 trimethylation during early embryonic development. PLoS Genet 10(8):e1004507.

35. Manna S, et al. (2015) Histone H3 Lysine 27 demethylases Jmjd3 and Utx are required for T-cell differentiation. Nat Commun 6:8152.

36. Cribbs A, et al. (2018) Inhibition of histone H3K27 demethylases selectively modulates inflammatory phenotypes of natural killer cells. J Biol Chem 293(7):2422–2437.

37. Kruidenier L, et al. (2012) A selective jumonji H3K27 demethylase inhibitor modulates the proinflammatory macrophage response. Nature 488(7411):404–408.

38. Heinemann B, et al. (2014) Inhibition of demethylases by GSK-J1/J4. Nature 514(7520):E1–2.

39. Satoh T, et al. (2010) The Jmjd3-Irf4 axis regulates M2 macrophage polarization and host responses against helminth infection. Nat Immunol 11(10):936–944.

40. Donas C, et al. (2016) The histone demethylase inhibitor GSK-J4 limits inflammation through the induction of a tolerogenic phenotype on DCs. J Autoimmun 75:105–117.

41. Sidrauski C, McGeachy AM, Ingolia NT, & Walter P (2015) The small molecule ISRIB reverses the effects of eIF2alpha phosphorylation on translation and stress granule assembly. Elife 4.

42. Cao Y, et al. (2019) ER stress-induced mediator C/EBP homologous protein thwarts effector T cell activity in tumors through T-bet repression. Nat Commun 10(1):1280.

43. Trapnell C, et al. (2014) The dynamics and regulators of cell fate decisions are revealed by pseudotemporal ordering of single cells. Nat Biotechnol 32(4):381–386.

44. Gerriets VA, et al. (2015) Metabolic programming and PDHK1 control CD4+ T cell subsets and inflammation. J Clin Invest 125(1):194–207.

45. Qureshi MA, Haynes CM, & Pellegrino MW (2017) The mitochondrial unfolded protein response: Signaling from the powerhouse. J Biol Chem 292(33):13500–13506.

46. Naruse C, et al. (2017) New insights into the role of Jmjd3 and Utx in axial skeletal formation in mice. FASEB J 31(6):2252–2266.

47. Burchfield JS, Li Q, Wang HY, & Wang RF (2015) JMJD3 as an epigenetic regulator in development and disease. Int J Biochem Cell Biol 67:148–157.

48. Welstead GG, et al. (2012) X-linked H3K27me3 demethylase Utx is required for embryonic development in a sex-specific manner. Proc Natl Acad Sci U S A 109(32):13004–13009.

49. Burgold T, et al. (2012) The H3K27 demethylase JMJD3 is required for maintenance of the embryonic respiratory neuronal network, neonatal breathing, and survival. Cell Rep 2(5):1244–1258.

50. Jiang Y, et al. (2018) Epigenetic activation during T helper 17 cell differentiation is mediated by Tripartite motif containing 28. Nat Commun 9(1):1424.

51. Miller SA, Mohn SE, & Weinmann AS (2010) Jmjd3 and UTX play a demethylase-independent role in chromatin remodeling to regulate T-box family member-dependent gene expression. Mol Cell 40(4):594–605.

52. Proserpio V & Mahata B (2016) Single-cell technologies to study the immune system. Immunology 147(2):133–140.

53. Gaublomme JT, et al. (2015) Single-Cell Genomics Unveils Critical Regulators of Th17 Cell Pathogenicity. Cell 163(6):1400–1412.

54. Wang C, et al. (2015) CD5L/AIM Regulates Lipid Biosynthesis and Restrains Th17 Cell Pathogenicity. Cell 163(6):1413–1427.

55. Wang R, et al. (2011) The transcription factor Myc controls metabolic reprogramming upon T lymphocyte activation. Immunity 35(6):871–882.

56. Fox CJ, Hammerman PS, & Thompson CB (2005) Fuel feeds function: energy metabolism and the T-cell response. Nat Rev Immunol 5(11):844–852.

57. Pearce EL & Pearce EJ (2013) Metabolic pathways in immune cell activation and quiescence. Immunity 38(4):633–643.

58. Donnelly RP, et al. (2014) mTORC1-dependent metabolic reprogramming is a prerequisite for NK cell effector function. J Immunol 193(9):4477–4484.

59. Nagai S, Kurebayashi Y, & Koyasu S (2013) Role of PI3K/Akt and mTOR complexes in Th17 cell differentiation. Ann N Y Acad Sci 1280:30–34.

60. Tameire F, et al. (2019) ATF4 couples MYC-dependent translational activity to bioenergetic demands during tumour progression. Nat Cell Biol 21(7):889–899.

61. Franchi L, et al. (2017) Inhibiting Oxidative Phosphorylation In Vivo Restrains Th17 Effector Responses and Ameliorates Murine Colitis. J Immunol 198(7):2735–2746.

62. Hough KP, Chisolm DA, & Weinmann AS (2015) Transcriptional regulation of T cell metabolism. Mol Immunol 68(2 Pt C):520–526.

63. Callegari S & Dennerlein S (2018) Sensing the Stress: A Role for the UPR(mt) and UPR(am) in the Quality Control of Mitochondria. Front Cell Dev Biol 6:31.

64. Quiros PM, et al. (2017) Multi-omics analysis identifies ATF4 as a key regulator of the mitochondrial stress response in mammals. J Cell Biol 216(7):2027–2045.

65. Merkwirth C, et al. (2016) Two Conserved Histone Demethylases Regulate Mitochondrial Stress-Induced Longevity. Cell 165(5):1209–1223.

66. Acosta-Rodriguez EV, Napolitani G, Lanzavecchia A, & Sallusto F (2007) Interleukins 1beta and 6 but not transforming growth factor-beta are essential for the differentiation of interleukin 17-producing human T helper cells. Nat Immunol 8(9):942–949.

67. Nakae S, Iwakura Y, Suto H, & Galli SJ (2007) Phenotypic differences between Th1 and Th17 cells and negative regulation of Th1 cell differentiation by IL-17. J Leukoc Biol 81(5):1258–1268.

68. Sornasse T, Larenas PV, Davis KA, de Vries JE, & Yssel H (1996) Differentiation and stability of T helper 1 and 2 cells derived from naive human neonatal CD4+ T cells, analyzed at the single-cell level. J Exp Med 184(2):473–483.

69. Stein CA, et al. (2010) Efficient gene silencing by delivery of locked nucleic acid antisense oligonucleotides, unassisted by transfection reagents. Nucleic Acids Res 38(1):e3.

70. Orlando DA, et al. (2014) Quantitative ChIP-Seq normalization reveals global modulation of the epigenome. Cell Rep 9(3):1163–1170.

71. Macosko EZ, et al. (2015) Highly Parallel Genome-wide Expression Profiling of Individual Cells Using Nanoliter Droplets. Cell 161(5):1202–1214.

72. Evans AM BB, Liu Q, Mitchell MW, Robinson RJ, Dai H, Stewart SJ, DeHaven CD and Miller LAD (2014) High Resolution Mass Spectrometry Improves Data Quantity and Quality as Compared to Unit Mass Resolution Mass Spectrometry in High-Throughput Profiling Metabolomics. Metabolomics 4(2).

73. Sims D, et al. (2014) CGAT: computational genomics analysis toolkit. Bioinformatics 30(9):1290–1291.

74. Cribbs A, et al. (2019) CGAT-core: a python framework for building scalable, reproducible computational biology workflows [version 1; peer review: 1 approved, 1 approved with reservations]. F1000Research 8(377).

75. Kim D, Langmead B, & Salzberg SL (2015) HISAT: a fast spliced aligner with low memory requirements. Nat Methods 12(4):357–360.

76. Liao Y, Smyth GK, & Shi W (2014) featureCounts: an efficient general purpose program for assigning sequence reads to genomic features. Bioinformatics 30(7):923–930.

77. Love MI, Huber W, & Anders S (2014) Moderated estimation of fold change and dispersion for RNA-seq data with DESeq2. Genome Biol 15(12):550.

78. Dobin A, et al. (2013) STAR: ultrafast universal RNA-seq aligner. Bioinformatics 29(1):15–21.

79. Langmead B, Trapnell C, Pop M, & Salzberg SL (2009) Ultrafast and memory-efficient alignment of short DNA sequences to the human genome. Genome Biol 10(3):R25.

80. Amir el AD, et al. (2013) viSNE enables visualization of high dimensional single-cell data and reveals phenotypic heterogeneity of leukemia. Nat Biotechnol 31(6):545–552.

