## Supplementary figures for "Histone H3K27me3 demethylases regulate human Th17 cell development and effector functions by impacting on metabolism"

**Table S1. Details of CyTOF antibodies used in this study.** Working volume and concentration represent the volume of the antibody added per 100 µl of staining buffer and the final concentration of antibody used for staining, respectively.

| Target | Tag | Working vol. or conc. |
|---|---|---|
| CD45 | 89Y | 1.5 µl |
| IdU | I 127 | 1 µl |
| CCR6 | Pr 141 | 1 µl |
| IL-4 | Nd 142 | 1 µl |
| CD127 | Nd 143 | 1.2 µl |
| CD38 | Nd 144 | 1.5 µl |
| CD4 | Nd 145 | 1.5 µl |
| CD8 | Nd 146 | 1 µg/ml |
| CD7 | SM 147 | 1.5 µl |
| IL-17a | Nd 148 | 1 µg/ml |
| CD200 | Sm 149 | 1 µg/ml |
| IL-22 | Nd 150 | 0.5 µl |
| TNF | Sm 152 | 1 µg/ml |
| CyclinB1 | Eu 153 | 2 µg/ml |
| CD3 | Sm 154 | 2 µg/ml |
| CD27 | Gd 155 | 1 µl |
| CD69 | Gd 156 | 1 µl |
| CD25 | Gd 158 | 1.5 µl |
| CD161 | Tb159 | 1 µl |
| TBET | Gd 160 | 1 µg/ml |
| Ki67 | Dy 161 | 2 µg/ml |
| FoxP3 | Dy 162 | 1.5 µl |
| CXCR3 | Dy 163 | 1 µl |
| CCR7 | Dy 164 | 2 µg/ml |
| IFNg | Ho 165 | 1 µl |
| pRb | Er 166 | 1.5 µl |
| GATA3 | Er 167 | 1 µl |
| ICOS | Er 168 | 1 µl |
| CD45RA | Tm 169 | 1.2 µl |
| HLA-DR | Er 170 | 1.5 µl |
| Granzyme B | Yb 171 | 1.2 µl |
| CD57 | Yb 172 | 0.5 µg/ml |
| CD137 | Yb 173 | 1.5 µl |
| PD-1 | Yb 174 | 1 µl |
| p-Histone | Lu 175 | 1 µl |
| CLA | Yb 176 | 1 µl |

**Table S2. Details of small molecular epigenetic probes used in this study.**

| Source Name | [Working] uM | Class/Target | SGC Global ID 1 |
|---|---|---|---|
| IOX1 (5-carboxy-8HQ) | 40 | Lysine demethylases - pan-2-OG | NCNNC183784a |
| Methylstat (Ester) | 2.5 | Histone demethylase | KDNMA000079 |
| (E)-JIB-04 | 0.05 | Histone demethylase - Pan JmjC | KDOOA011528a |
| KDOBA67 | 10 | Histone demethylase | |
| GSK J4 | 10 | Lysine demethylases - JMJD3, UTX, JARID1B | KDGOA00000(1-8) |
| GSK J5 (inactive) | 10 | Lysine demethylases - Negative control | KDGOA00000(1-8) |
| KDOBA67 | 10 | Histone demethylase | |
| GSK-LSD1 (irreversible) | 0.5 | Lysine demethylases - LSD1 | |
| (+)-JQ1 | 1 | Bromodomains - BRD2, BRD3, BRD4, BRDT (BET) | |
| (-)-JQ1 (inactive) | 1 | Bromodomains - Negative control | |
| PFI-1 | 5 | Bromodomains - BRD2, BRD3, BRD4, BRDT (BET) | BDF00004323 |
| I-BET | 1 | Bromodomains - BRD2/3/4 | |
| Bromosporine | 1 | Bromodomains - pan-Bromodomain | BDOTC000133 |
| CBP/BRD4 (0383) | 5 | Bromodomains - CBP, BRD4(1) | BDOIA000383 |
| SGC-CBP30 | 1 | Bromodomains - CREBBP, EP300 | BDOIA000518a |
| I-CBP112 | 1 | Bromodomains - CREBBP, EP300 | BDOBO000038a |
| RVX-208 | 5 | Bromodomains - BRD2, BRD3, BRD4, BRDT (BET, BD2) | BDOOA011571a |
| GSK2801 | 1 | Bromodomains - BAZ2A, BAZ2B | BDG00021335 |
| CXD101 | 1 | HDAC - | |
| Valproic acid | 1000 | HDAC - aliphatic acid compounds | |
| Entinostat | 0.5 | HDAC - ortho-amino anilides | |
| Trichostatin A | 0.5 | HDAC - hydroxamic acids - Class I & II | SDOOA001122 |
| CI-994 | 1 | HDAC - 1,2,3,(8) | |
| Rucaparib | 10 | Poly ADP ribose polymerase (PARP) | |
| SAHA | 2.5 | HDAC - hydroxamic acids | HDTTA000395 |
| Belinostat | 5 | HDAC - hydroxamic acids | |
| SRT1720 | 1 | HDAC - SIRT1 (indirect?) activator | |
| EX 527 | 1 | HDAC - SIRT1 | |

| | | | |
|---|---|---|---|
| UNC1999 | 1 | Histone methyltransferase - EZH2 | |
| 5-Azacitidine | 10 | DNA methyltransferase (DNMT) - | NOT FOUND |
| 5-Azadeoxycitidine | 5 | DNA methyltransferase (DNMT) - DNMT1/3 | NOT FOUND |
| C646 | 1 | Histone acetyltransferase (HAT) p300/CBP | |
| Chaetocin | 0.05 | Histone methyltransferase - SUV39H1 | NOT FOUND |
| UNC1215 | 5 | Methyl Lysine Binder - L3MBTL3 | MBU00001215 |
| GSK343 | 3 | Histone methyltransferase - EZH2 | |
| PFI-2 | 2 | Histone methyltransferase - SETD7 | |
| UNC0638 | 1 | Histone methyltransferase - G9a, GLP | MTUFP000001a |
| UNC0642 | 1 | Histone methyltransferase - G9a, GLP | |
| A-366 | 2 | Histone methyltransferase - G9a, GLP | |
| SGC0946 | 7.5 | Histone methyltransferase - DOT1L | MTT00000946 |
| IOX2 | 10 | Prolyl-Hydroxylases - PHD2 (EGLN1) | KDOOA010881 |
| SMARCA | 2.5 | Bromodomains - SMARCA, PB1 | BDF00008718 |
| Olaparib | 1 | Poly ADP ribose polymerase (PARP) | NOT FOUND |
| K00135 | 1 | Kinase inhibitor - ATP competitive - PIM | PKOOA000135 |

**Table S3. Details of primers used to detect mRNA levels for RT-qPCR in this study.**

| Gene | Forward primer | Reverse primer |
|---|---|---|
| IFNG | TCGGTAACTGACTTGAATGTCCA | TCGCTTCCCTGTTTTAGCTGC |
| ATF4 | ATGACCGAAATGAGCTTCCTG | GCTGGAGAACCCATGAGGT |
| ATF3 | CCTCTGCGCTGGAATCAGTC | TTCTTTCTCGTCGCCTCTTTTT |
| DDIT3 | GGAAACAGAGTGGTCATTCCC | CTGCTTGAGCCGTTCATTCTC |
| INHBE | ATCTTCCGATGGGGACCAAG | AGAGTTAAGGTATGCCAGCCC |
| KDM6B | CACCCCAGCAAACCATATTATGC | CACACAGCCATGCAGGGATT |
| KDM6A | TTCCTCGGAAGGTGCTATTCA | GAGGCTGGTTGCAGGATTCA |
| KDM5B | CCATAGCCGAGCAGACTGG | GGATACGTGGCGTAAAATGAAGT |
| IL17A | TCCCACGAAATCCAGGATGC | GGATGTTCAGGTTGACCATCAC |
| IL17F | GCTGTCGATATTGGGGCTTG | GGAAACGCGCTGGTTTTCAT |
| RORC | GTGGGGACAAGTCGTCTGG | AGTGCTGGCATCGGTTTCG |

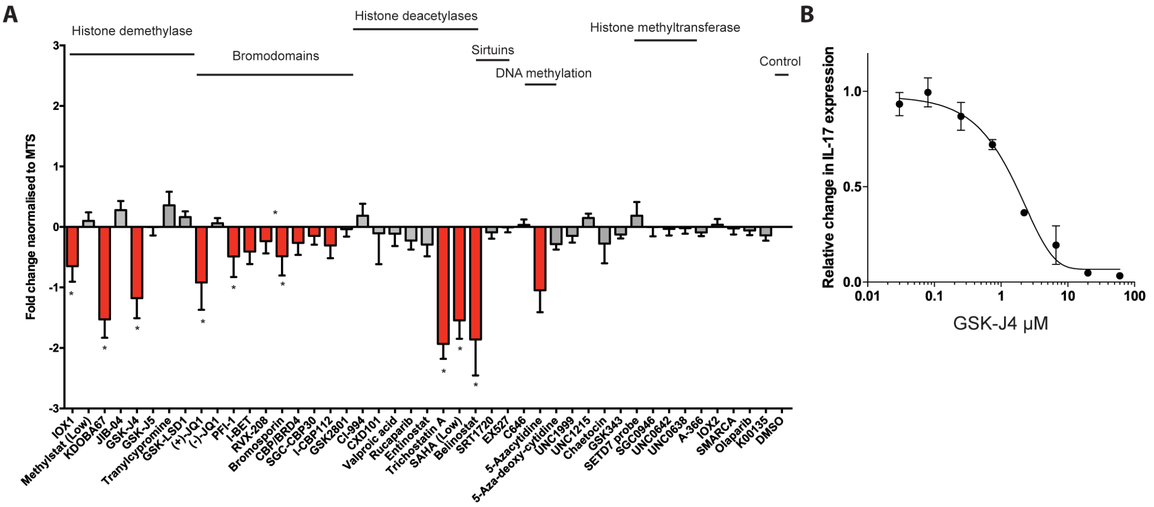


**Figure S1**: Small molecule epigenetic compound screening reveals BET bromodomains, Histone deacetylases and Jumonji histone demethylases as regulators of IL17 production. (A) Relative expression of IL-17 expression from Th17 differentiated cells, as measured by ELISA and normalised to MTS assay. (B) Plot showing the secretion of IL-17 from Th17 differentiated cells, as measured by ELISA, over a range of GSK-J4 concentrations. Data are mean ± S.D. P values were calculated using Kruskal-Wallis test. *P < 0.05.

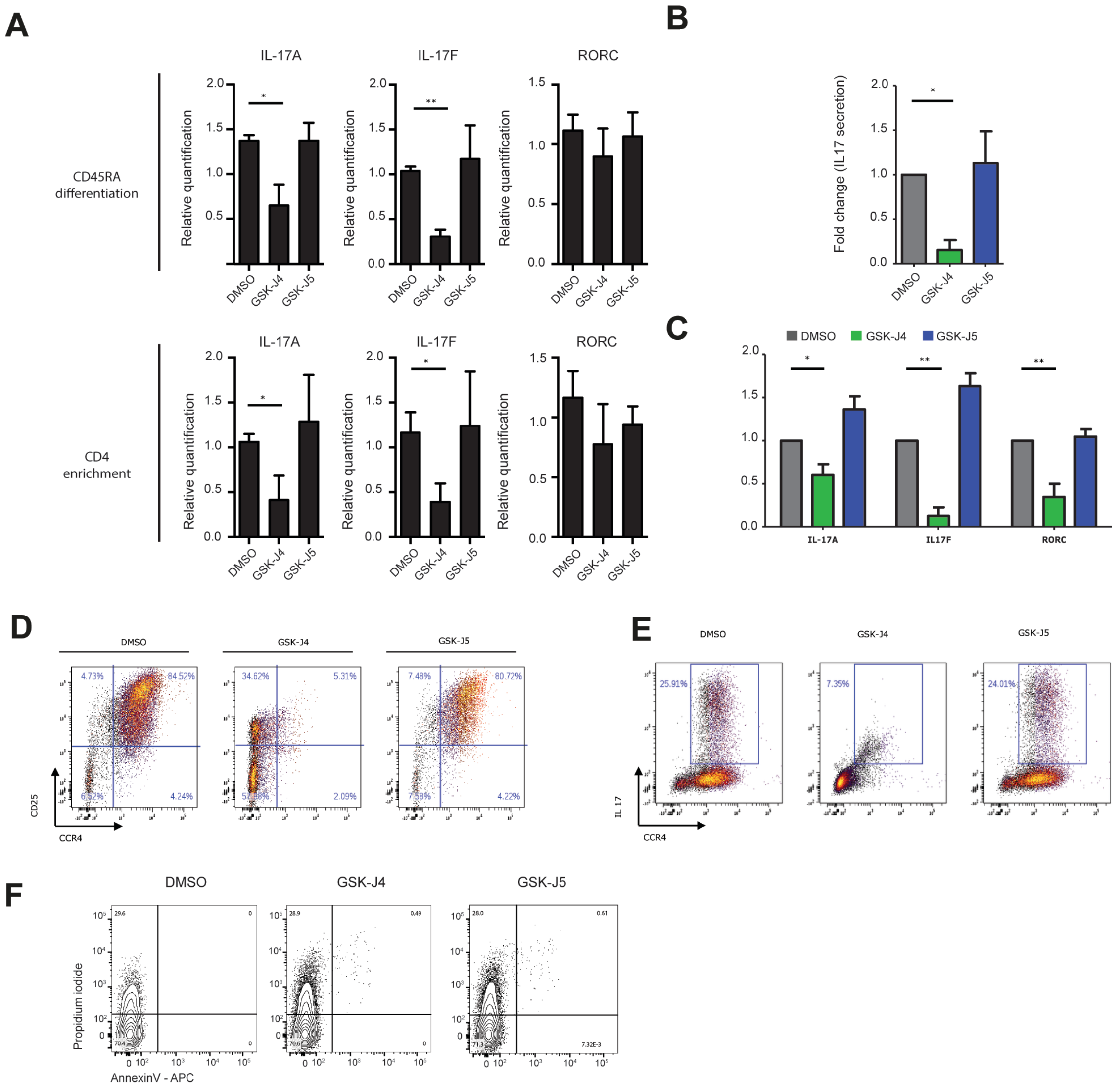


**Figure S2**: KDM6B/KDM6A inhibition by GSK-J4 reduces pro-inflammatory function in both enriched and differentiated Th17 cells. (A) RT-qPCR showing the quantification of IL-17A, IL-17F and RORC following 24 hours of GSK-J4 treatment. Top panel shows the effect of GSK-J4 on Th17 differentiated from CD45RA$^{+}$ naïve cells and bottom panel shows the effect of GSK-J4 on Th17 cells enriched from CD4$^{+}$ cells. (B) IL17 expression measured by ELISA, following addition of GSK-J4 over the course of the 7 days differentiation protocol. (C) RT-qPCR showing the quantification of IL-17A, IL-17F and RORC with GSK-J4 treatment added before and during the full differentiation protocol. (D) Flow cytometric analysis of CD25 and CCR4 staining in Th17 differentiated cells that have been treated with GSK-J4 during the full 7-day differentiation protocol. (E) Flow cytometric analysis of IL17 and CCR4 staining in Th17 cells following GSK-J4 treatment during the full 7-day differentiation protocol. (F) PI and Anexin-V staining of Th17 cells at day 7 following treatment with GSK-J4 added before and during the full differentiation protocol. Data are mean ± S.D. P values were calculated using Kruskal-Wallis test. **P < 0.01, *P < 0.05.

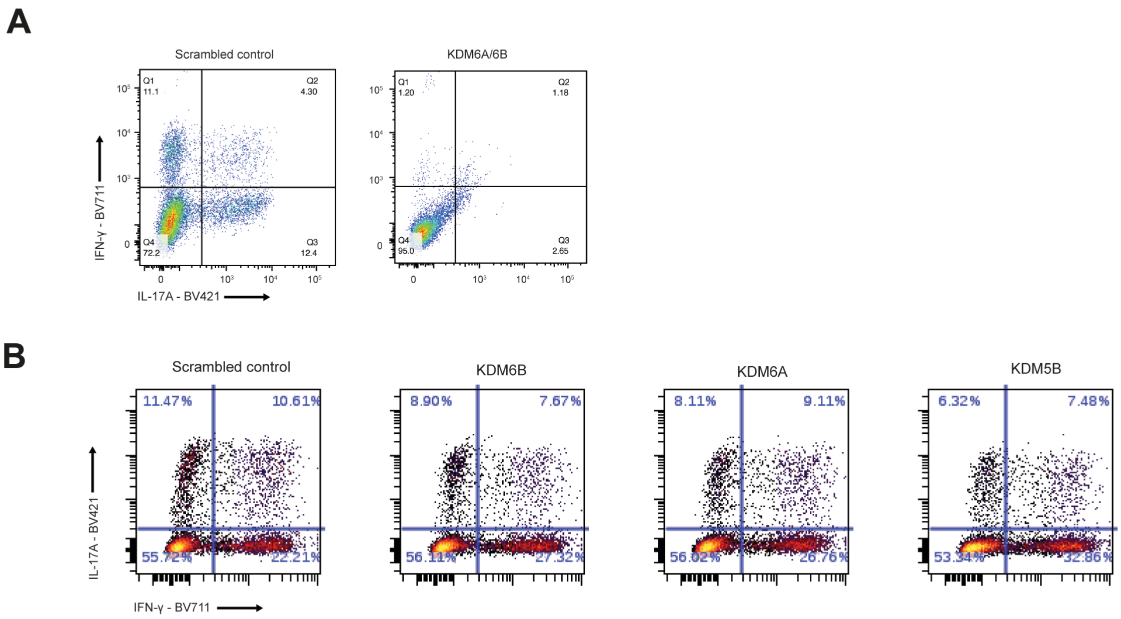


Fig. S3: KDM6B and KDM6A knockdown on their own only mildly inhibits cytokine production in Th17 differentiated cells. (A) LNA dual knockdown of KDM6A/KDM6B and scrambled control was performed over 7 days of Th17 differentiation from naïve precursors. Cells were stimulated with PMA/ionomycin for 4 hours then IL17 and IFN-γ was measured using flow cytometry. (B) LNA knockdown of KDM6B, KDM5A and KDM5B was performed over 7 days of Th17 differentiation from naïve precursors and the expression of IL-17A and IFN-γ was measured using flow cytometry. Data are representative of 4 individual experiments.

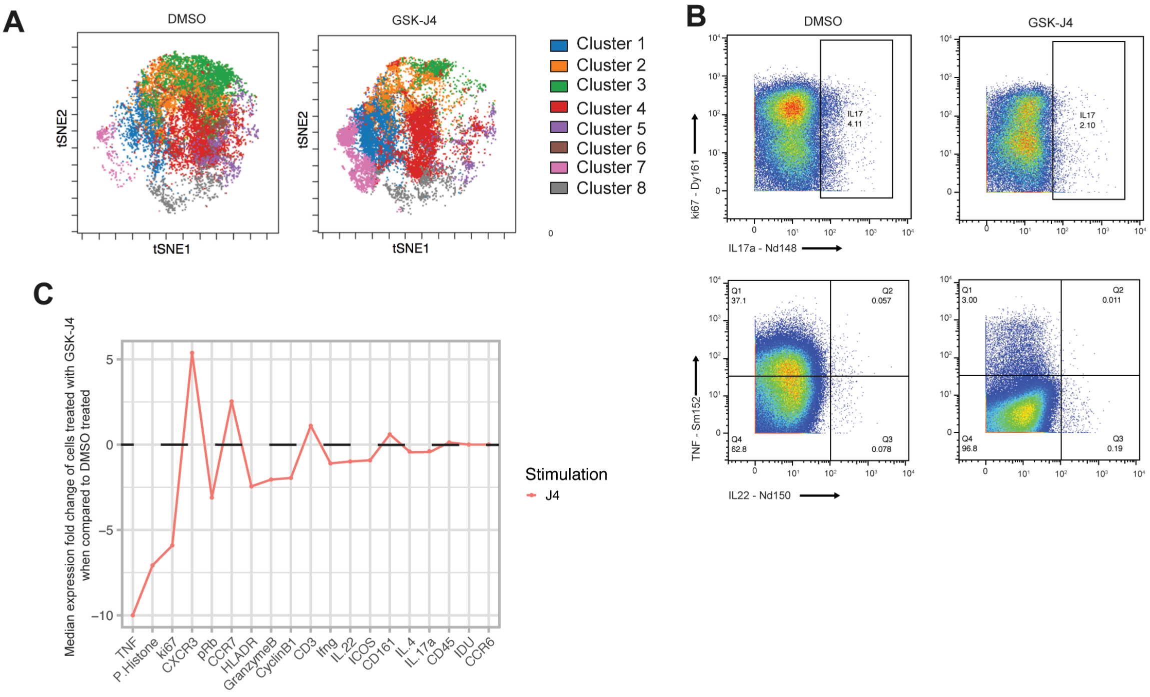


**Figure S4**: CyTOF analysis reveals significantly altered cell cycle response to GSK-J4. (A) tSNE plot showing FlowSOM clustering analysis of Th17 differentiated cells treated with either DMSO or GSK-J4 for 24 hours. (B) CyTOF analysis of IL-22 and IL17A cytokine expression of Th17 differentiated cells following 24 hours of DMSO or GSK-J4 treatment. (C) Relative fold change in median expression values for GSK-J4 when compared to DMSO. (D) A plot showing the relative expression proportion for cell cycle markers displayed as a radial visual plot. CyTOF plots are representative of 3 individual donors.

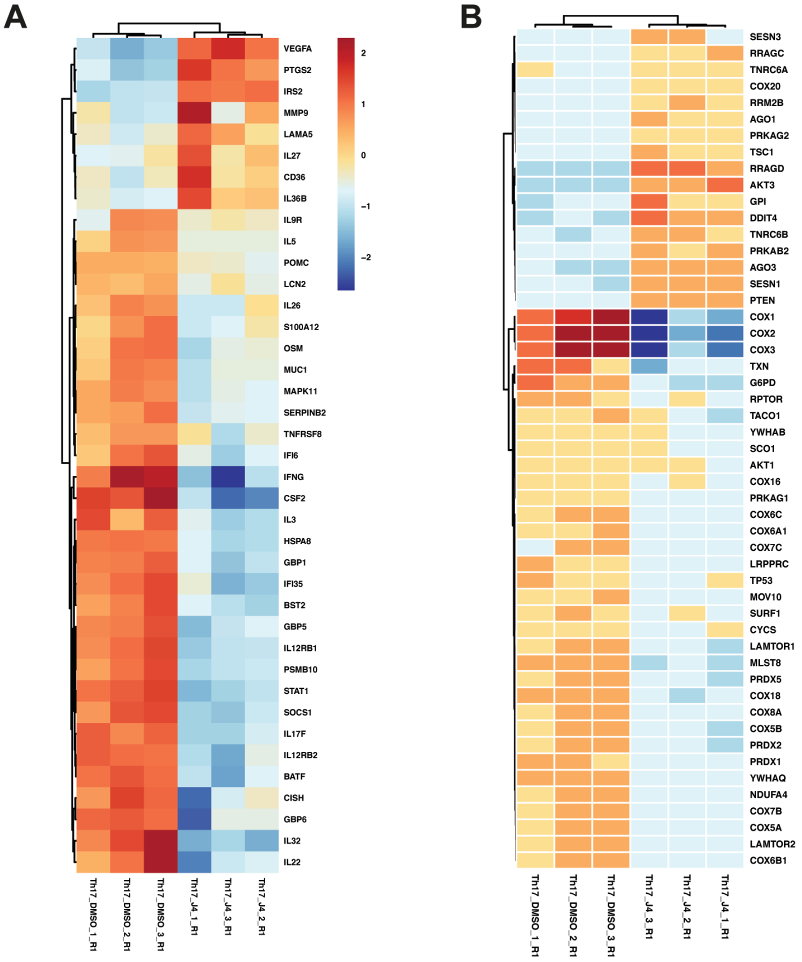


**Figure S5**: RNA-seq highlights downregulation of interleukin signalling, the induction of an ATF4 transcription factor family response. (A) Heatmap showing genes associated with interleukin signalling. (B) Heatmap showing genes associated with the pathway TP53 regulates metabolic gene expression

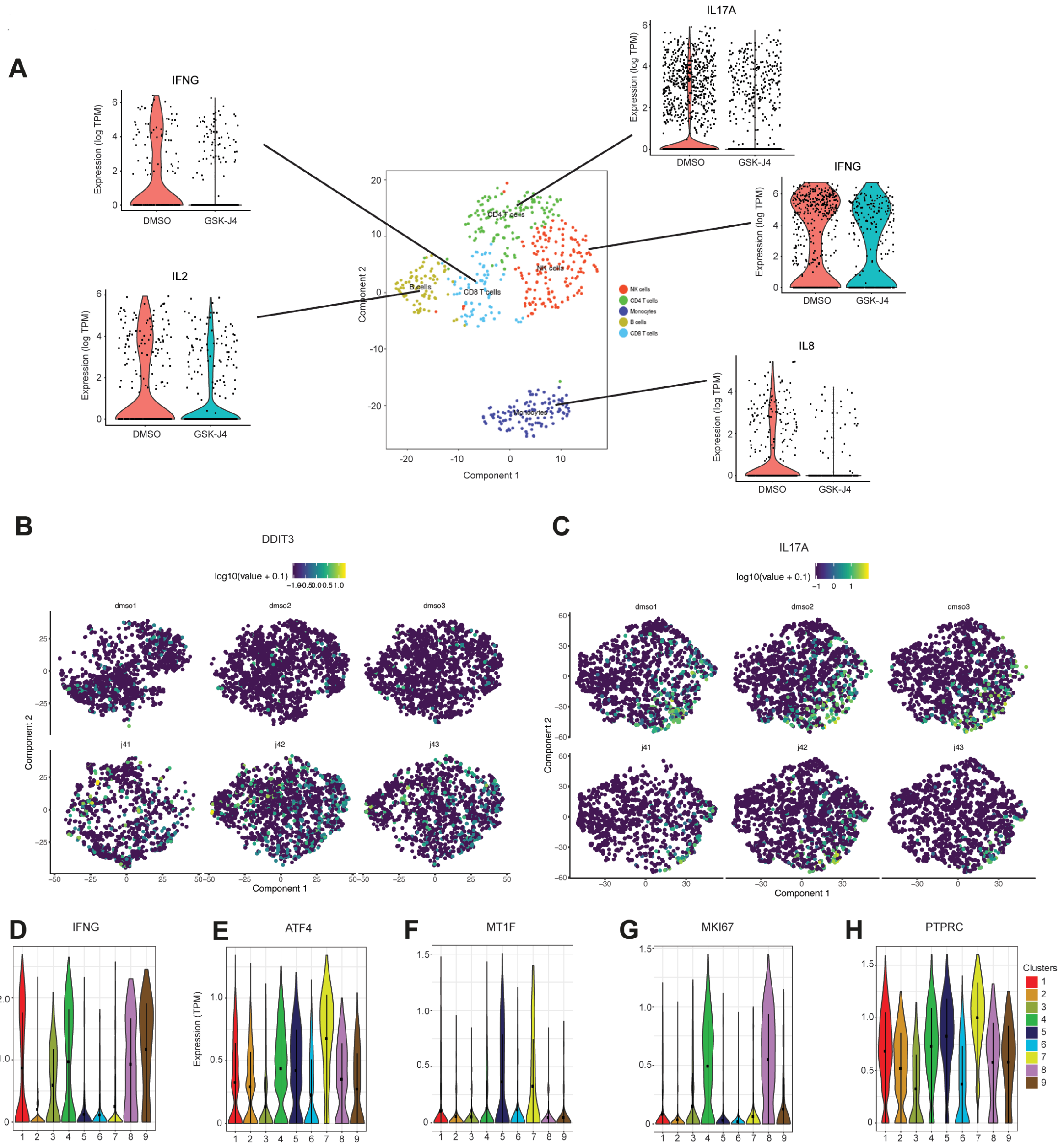


**Figure 6S**: Single cell sequencing reveals significantly altered heterogeneity in response to GSK-J4 treatment. (A) A PBMC sample was isolated from healthy individual and cultured in the presence of DMSO or GSK-J4 for 24 hours then scRNA-seq was performed. tSNE clustering highlights the different immune populations identified by marker expression. CD8 T cells were identified as CD3$^+$CD8$^+$, CD4 T cells CD3$^+$CD4$^+$, B cells as CD20$^+$, NK cells as CD3$^-$CD56$^+$ and monocytes as CD14$^+$. The top differentially expressed cytokine between DMSO and GSK-J4 is shown for each population. (B) tSNE plots of Th17 differentiated cells from AS donors showing the expression of DDIT3 in 3 independent donors. (C) tSNE plots of Th17 differentiated cells from AS donors showing the expression of IL17A in 3 independent donors. The expression of (D) ATF4, (E) IFNG, (F) MT1F, (G) MKI67 and (H) PTPRC in each of the Th17 clusters identified in Figure 5.

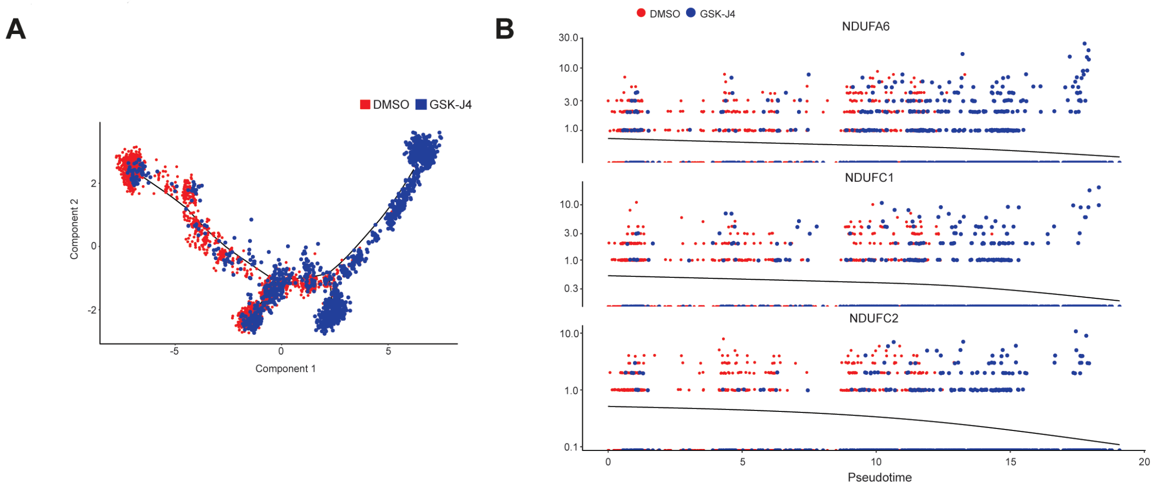


**Figure S7**: Pseudotime analysis reveals downregulation of genes associated with the TCA cycle following GSK-J4 treatment. TCA related genes NDUFA6, NDUFC1 and NDUFC2 are plotted across pseudotime.

### FIGURE S8

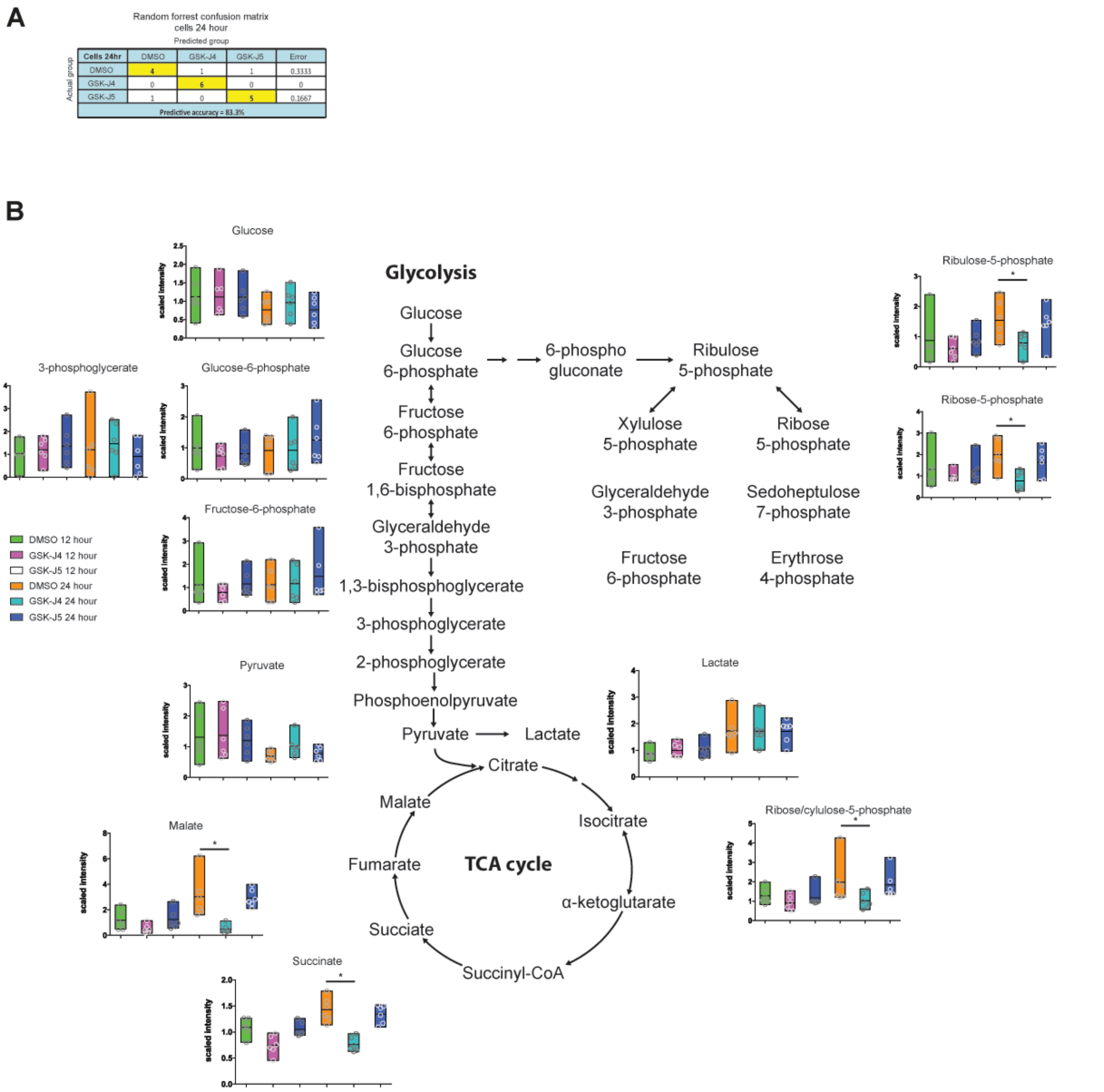


**Figure S8**: The glycolysis pathway showing switch to anaerobic glycolysis. (A) Random forest confusion matrices showing the correlation between drug treatments. (B) The glycolysis pathway showing the all metabolites measured during mass spectrometry. Values are means ± SEM; P values were calculated using Welch's t-test. **P* < 0.05; **P < 0.01; n=5

### FIGURE S9

A

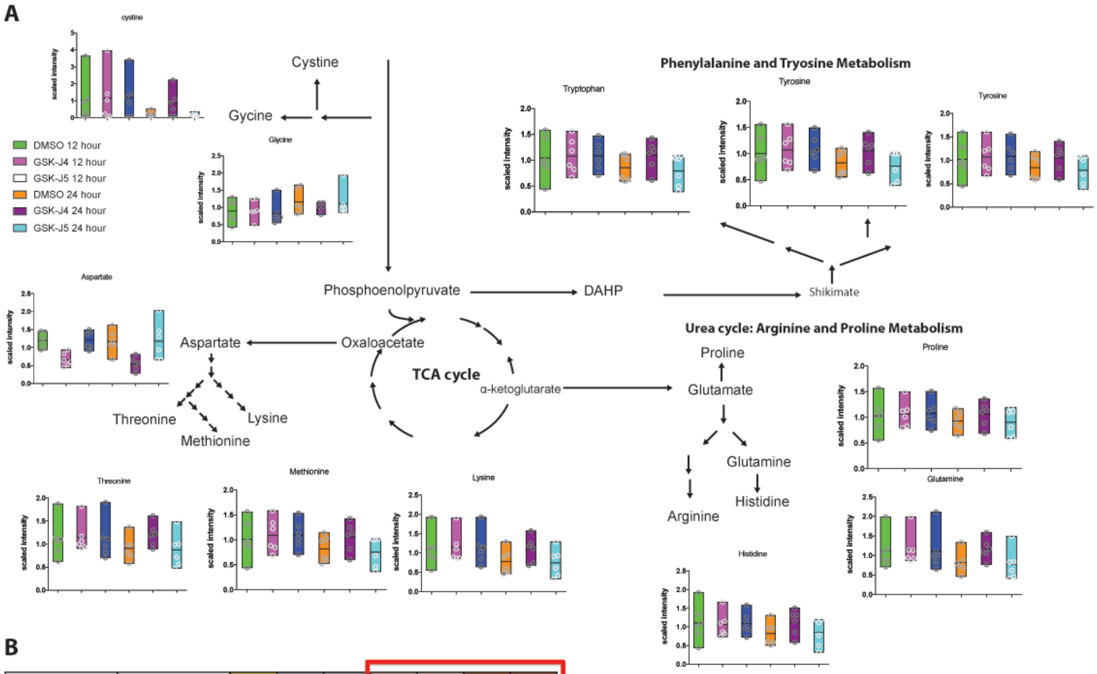


B

| Sub Pathway | Biochemical Name | BL POST DIFF BL PRE DIFF | DMSO 12hr BL POST DIFF | DMSO 24hr BL POST DIF | J4 12HR DMSO 12HR | J5 12HR DMSO 12HR | J4 24HR DMSO 24HR | J5 24HR DMSO 24HR |
|---|---|---|---|---|---|---|---|---|
| Purine Metabolism, Xanthine/Inosine containing (*Cells*) | inosine | 4.51 | 0.69 | 0.82 | 0.66 | 0.88 | 0.41 | 0.53 |
| | hypoxanthine | 3.36 | 0.77 | 0.92 | 0.78 | 0.79 | 0.6 | 0.85 |
| | 2'-deoxyinosine | 4.45 | 1.64 | 1.29 | 0.5 | 0.41 | 0.32 | 0.56 |
| | urate | 0.44 | 1 | 1 | 1 | 1 | 1 | 1 |
| Purine Metabolism, Adenine containing (*Cells*) | ADP | 0.28 | 0.73 | 0.83 | 1.04 | 0.65 | 0.65 | 1.47 |
| | AMP | 1.71 | 0.77 | 1.02 | 0.76 | 0.94 | 0.67 | 0.9 |
| | 2'-AMP | 1.09 | 0.65 | 0.64 | 0.74 | 0.79 | 0.61 | 0.73 |
| | cAMP | 1.01 | 0.72 | 0.57 | 0.64 | 0.9 | 0.75 | 0.91 |
| | adenylosuccinate | 1.6 | 2.19 | 9.79 | 0.51 | 1.24 | 0.11 | 0.71 |
| | adenosine | 1.46 | 0.57 | 0.84 | 1.11 | 0.96 | 0.73 | 1.25 |
| | adenine | 0.9 | 1.37 | 1.28 | 1.06 | 0.78 | 0.84 | 0.98 |
| | N1-methyladenosine | 0.93 | 0.4 | 0.54 | 2.49 | 4.47 | 2.27 | 2.9 |
| | N6-carbamoylthreonyladenosine | 1.56 | 0.43 | 0.41 | 1.07 | 1.4 | 1.85 | 1.45 |
| | 2'-deoxyadenosine 5'-monophosphate | 7.37 | 0.98 | 3.36 | 0.71 | 0.97 | 0.39 | 0.92 |
| Pyrimidine Metabolism, Orotate containing (*Cells*) | N-carbamoylaspartate | 1.07 | 2.12 | 116.47 | 38.18 | 62.68 | 0.43 | 1.64 |
| | orotate | 1.52 | 2.48 | 31.58 | 6.08 | 8.46 | 0.45 | 1.09 |
| | orotidine | 1.4 | 1.6 | 3.67 | 0.57 | 0.84 | 0.21 | 1.03 |
| Pyrimidine Metabolism, Uracil containing (*Cells*) | uridine 5'-diphosphate (UDP) | 0.53 | 1.6 | 1.45 | 0.88 | 0.63 | 0.82 | 2.11 |
| | uridine 5'-monophosphate | 4.85 | 1.73 | 2.57 | 0.49 | 0.71 | 0.43 | 1.01 |
| | uridine | 3.33 | 1.01 | 1.12 | 0.72 | 0.8 | 0.59 | 0.74 |
| | pseudouridine | 0.73 | 0.46 | 0.92 | 1.26 | 1.76 | 0.98 | 1.08 |
| | 5-methyluridine (ribothymidine) | 2.39 | 0.62 | 0.85 | 0.8 | 1.09 | 0.8 | 0.74 |
| | beta-alanine | 1.46 | 1.12 | 0.75 | 0.8 | 0.77 | 0.84 | 1.13 |
| Pyrimidine Metabolism, Cytidine containing (*Cells*) | cytidine diphosphate | 0.81 | 2.39 | 1.68 | 1.04 | 0.57 | 1.1 | 2.26 |
| | 5'-CMP | 3.8 | 1.34 | 1.68 | 0.7 | 0.88 | 0.71 | 0.91 |
| | cytidine | 14.15 | 1.01 | 1.19 | 0.73 | 0.8 | 0.62 | 0.7 |
| | 2'-deoxycytidine | 1 | 1 | 3.04 | 1 | 1 | 0.33 | 0.88 |
| Pyrimidine Metabolism, Thymine containing (*Cells*) | thymidine 5'-monophosphate | 9.45 | 1.13 | 3.14 | 0.62 | 0.86 | 0.49 | 0.9 |
| | thymidine | 0.9 | 1.32 | 0.77 | 0.65 | 0.65 | 0.78 | 0.74 |
| | thymine | 1.66 | 1.45 | 1.11 | 0.67 | 0.5 | 0.86 | 0.98 |

**Figure S9**: Significant downregulation of multiple nucleotide metabolites. (A) The TCA cycle showing the amino acid products measured during mass spectrometry following DMSO, GSK-J4 and GSK-J5 treatment of Th17 differentiated cells for 24 hours. Values are means ± SEM; **P* < 0.05; **P < 0.01; ***P < 0.001; n=5 (B) Table showing the downregulation of a number of nucleotide metabolites following 24 hours of GSK-J4 treatment. Values are means ± SEM; **P* < 0.05
